# FAF1 cofactor enhances UFD1/NPL4-p97 unfolding efficiency across ubiquitin chain lengths independent of SUMO2

**DOI:** 10.64898/2026.07.31.742138

**Authors:** Abigail A. Lemmon, Christopher D. Lima

## Abstract

The AAA+ protein p97/VCP and its cofactor UFD1/NPL4 interact with and unfold ubiquitinated proteins to promote disaggregation and unfolding for recycling or to prepare substrates for proteasomal degradation. The cofactor Fas-associated factor 1 (FAF1) is suggested to reduce the length of ubiquitin chain required for substrate unfolding by UFD1/NPL4-p97 and to interact with SUMO. Here, we employ in vitro reconstitution of UFD1/NPL4-p97 and FAF1/UFD1/NPL4-p97 complexes and fluorescent substrates modified with SUMO2-polyubiquitin hybrid or polyubiquitin-only chains of varying lengths to assess initial rates of unfolding. These assays reveal that FAF1 enhances initial rates of unfolding relative to UFD1/NPL4-p97 in a manner that is independent of SUMO2 and semi-dependent on ubiquitin chain length. Unlike preferences observed for yeast Ufd1/Npl4-Cdc48, these data suggest that the FAF1 cofactor does not contribute to preferential unfolding of the SUMO2-polyubiquitin substrates tested. Further dissection of FAF1 reveals that it significantly increases the rate of unfolding for all ubiquitin chain lengths tested with its greatest differential impact observed when unfolding chains with four to ten ubiquitin molecules that are considered physiologically relevant. Using cryoEM we resolve a series of reconstructions that reveal FAF1/UFD1/NPL4-p97 bound to substrate in non-translocating and translocating states. Observed interactions between a helix of FAF1 and UFD1 throughout the unfolding process are consistent with AlphaFold models and recent reports suggesting that FAF1 may stabilize interactions between UFD1, NPL4, and p97 to promote substrate engagement and unfolding.

## Introduction

Post-translational modification by ubiquitin and ubiquitin-like proteins such as SUMO (Small Ubiquitin-like Modifier) is a key mechanism to regulate the quality and levels of protein. Conjugation of ubiquitin and SUMO occurs primarily on lysine residues of target proteins and is catalyzed via a three-enzyme cascade consisting of E1 activating enzymes, E2 conjugating enzymes, and E3 protein ligases (1, 2). Further modification by ligases and proteases can generate a diversity of polyubiquitin linkage types and chain topologies in addition to substrates modified by more than one type of ubiquitin-like protein (3). Various factors can specifically recognize different chain topologies and linkage types to guide modified proteins toward specific fates (4–6). Proteins conjugated with a polyubiquitin chain linked via lysine 48 of ubiquitin (K48 chains) are canonically targeted to the proteasome for subsequent degradation (7–9). A subset of SUMOylated proteins are conjugated with polyubiquitin via SUMO-polyubiquitin hybrid chains after modification by SUMO-targeted ubiquitin ligases (STUbLs) (10). These substrates are then targeted for disaggregation or to the proteasome for degradation (11). For proteins to be processed by the proteasome, substrates must have an unstructured initiation site, which frequently requires further processing, disaggregation, or extraction prior to degradation (12).

The AAA+ protein p97 (VCP, Cdc48 in yeast) is responsible for disaggregating or unfolding proteins to prepare them for recycling or degradation by the 26S proteasome (13–15). Unfolding by p97 is involved in cellular processes including endoplasmic reticulum associated degradation (ERAD) (16, 17), membrane fusion (18, 19), and nuclear activities (20–25). The importance of p97 is highlighted by diseases defined by p97 mutation and misfunction, particularly neurodegenerative disorders (26–28). p97 is a homo-hexameric ring that includes two ATPase D1 and D2 domains, and an N terminal domain (29) which is the primary binding site for cofactors (30, 31). A wide array of cofactors are responsible for regulating the functions of p97 (32, 33). The best characterized cofactor is the heterodimer UFD1/NPL4 (UN) which is evolutionarily conserved and preferentially binds to and unfolds substrates modified by K48-linked ubiquitin chains (34, 35). UFD1 contributes to ubiquitin chain topology specificity (36) and binds to p97 via SHP motifs while NPL4 binds to ubiquitin chains, unfolds ubiquitin, and interacts with p97 via zinc fingers and a ubiquitin regulatory X (UBX)-like domain (UBXL) (37, 38). UFD1/NPL4-p97 (UN-p97) initiates unfolding of a ubiquitinated substrate in an ATP-independent manner via binding of the unfolded initiator ubiquitin to a hydrophobic groove in NPL4 which then allows for insertion into the central pore of p97 (39). p97 then utilizes ATP hydrolysis to generate conformational changes within the hexameric ring to pull the unfolded substrate peptide through the pore using a hand over hand mechanism shared among other AAA+ ATPases (40). While unfolding can initiate on any ubiquitin within a K48 chain, productive unfolding proceeds by unfolding the ubiquitin that is N-terminal and proximal to the substrate (41). Chain topology, linkage type, and chain length are therefore major determinants in the activity of p97 complexes.

We previously demonstrated that a C-terminal SUMO interaction motif (SIM) in yeast Ufd1 results in preferential unfolding of substrates modified by SUMO (Smt3)-polyubiquitin hybrid chains (42). The SIM of Ufd1 is conserved in fission and budding yeasts, but is not conserved in mammals, suggesting that human UN-p97 may not interact with SUMO-polyubiquitin chains in similar manner to yeast UN-Cdc48. As substrates containing SUMO-polyubiquitin hybrid chains and SUMO-polyubiquitin readers are present in human (43), we sought to determine additional cofactors that enable recognition of SUMO-polyubiquitin hybrid chains by p97. We focused on Fas-associated factor 1 (FAF1) as it has been suggested to interact with SUMO1 in a yeast two-hybrid screen (44) and the *Caenorhabditis elegans* homolog interacts with SUMO *in vitro* (45). Additionally, upon inhibition of a SUMO-targeted deubiquitinase, there is increased colocalization of FAF1-bound p97 with SUMOylated proteins (45), suggesting that FAF1 may be responsible for clearance of SUMO-polyubiquitin conjugated proteins. A direct role for FAF1 in unfolding SUMO-polyubiquitin hybrid chains remains less clear, but its role in ubiquitin-targeted unfolding including ERAD (46) has been studied. FAF1 interacts with p97 via a UBX domain (47) and has a ubiquitin-associated (UBA) domain which interacts with ubiquitin (48) with a preference for K48 linked chains (49). FAF1 interacts with UN-p97 to form a complex of FAF1/UFD1/NPL4-p97 (FUN-p97). Recent studies revealed that UN-p97 requires longer ubiquitin chains for productive unfolding compared to yeast UN-Cdc48 complex, and that addition of UBX-containing proteins such as FAF1 reduce the ‘ubiquitin threshold’ to one comparable to the yeast complex (50). In addition to its potential interaction with SUMO-polyubiquitin hybrid chains, the mechanism by which FAF1 enhances unfolding remained unclear until recent studies suggesting that a helical domain of FAF1 may stabilize UFD1 in a FUN-p97 complex (51–56).

Here we reconstitute the FUN-p97 and UN-p97 complexes unfolding fluorescent substrates modified with SUMO-polyubiquitin hybrid or polyubiquitin-only chains with varying ubiquitin chain lengths to assess the effects of SUMO and chain length on unfolding by respective complexes. We determine that neither SUMO2 nor polySUMO2 enhance unfolding rates at any ubiquitin chain length, suggesting that FAF1 does not preferentially unfold SUMO2-polyubiquitin hybrid chains of the topologies tested. We show that FUN-p97 unfolds a polyubiquitinated substrate more efficiently than UN-p97 at all ubiquitin chain lengths tested, with greatest effects observed for substrates modified by polyubiquitin chains containing 4-10 ubiquitins. We also observe that the UBA domain of FAF1 is dispensable for unfolding, and that the FAF1 helical domain and UBX domain are sufficient to increase initial rates of unfolding for all ubiquitin chain lengths tested, extending recent observations reported by others (51–56). Using single particle analysis and cryogenic electron microscopy (cryo-EM) we report reconstructions of FUN-p97 complexes in the presence of a polyubiquitin substrate that capture the complex in multiple states representing substrate-bound complex prior to initiating unfolding, substrate-bound complexes after unfolding initiation but prior to threading of substrate into the p97 pore, and complexes actively unfolding substrate, as well as a complex in the absence of substrate.

Reconstructions are consistent with a single FAF1 bound to p97 via its UBX domain and binding to and potentially stabilizing UFD1 via its helical domain. Notably, the UT3 domain of UFD1 is positioned near ubiquitin binding sites of NPL4 in the presence of FAF1. Weaker density on the adjacent p97 N-domain suggests that the UBX domain of a second FAF1 may be bound to p97, however most classes exhibit weak or no densities for the helical domain, suggesting that a single FAF1 is responsible for UFD1 interaction.

## RESULTS

### Biochemical reconstitution of human SUMO-polyubiquitin modified substrates

Substrates containing SUMO-polyubiquitin (polyUb) hybrid chains were biochemically generated based on published protocols (42) utilizing a photo-convertible fluorescent substrate mEos (14, 57) with a linear fusion of human SUMO2 and human ubiquitin (SUMO2-Ub-mEos). The single ubiquitin within the SUMO2-Ub-mEos substrate was conjugated with ubiquitin to form ubiquitin chains in assays containing E1 *Schizosaccharomyces pombe (sp)* Uba1, K48-specific chain-elongating E2 *Homo sapiens (hs)* UBE2K (58), and STUbL *sp*Slx8-Rfp2 (59) (Fig. 1A, Fig. S1A). Slx8-Rfp2 was used to conjugate ubiquitin to the SUMO2-Ub-mEos fusion as it requires a single SUMO moiety for activity thus bypassing human RNF4’s dependence on polySUMO (60). Consistent with the Ub within SUMO-Ub-mEos being modified at position K48, ubiquitin chain extension was diminished for a SUMO2-UbK48R-mEos fusion (Fig. S1B). Further, cleavage by ubiquitin-specific peptidase 2 (Usp2) (61) led to collapse of chains into ubiquitin, SUMO2-Ub, and mEos bands, and treatment with both Usp2 and SUMO-specific peptidase 1 (SENP1) lead to collapse into ubiquitin, SUMO2, and mEos bands (Fig. S1C). After modification, substrates were separated using size exclusion chromatography to separate substrates containing polyUb chains of varying lengths with 0-4 Ub (low, L), 2-5 Ub (medium, M), 4-10 Ub (high, H), and >6 Ub (very high, VH) (Fig. 1A, Fig. S1D). We also generated a polySUMO2 substrate containing three SUMO2 moieties fused end to end to ubiquitin and mEos (3xSUMO2-Ub-mEos) which was then modified with a polyUb chain containing 4-10 ubiquitin proteins (Fig. 1A, Fig. S2A). SUMO2-polyUb-mEos and 3xSUMO2-Ub^H^-mEos substrates were photoconverted as described previously (14, 42) to generate a bipartite substrate that loses fluorescence when unfolded. (poly)SUMO-polyUb-mEos photoconverted substrates are referred to as SUMO2-Ub^X^ or 3xSUMO2-Ub^X^ with superscript denoting ubiquitin chain length (L, M, H, or VH).

**Figure 1.**
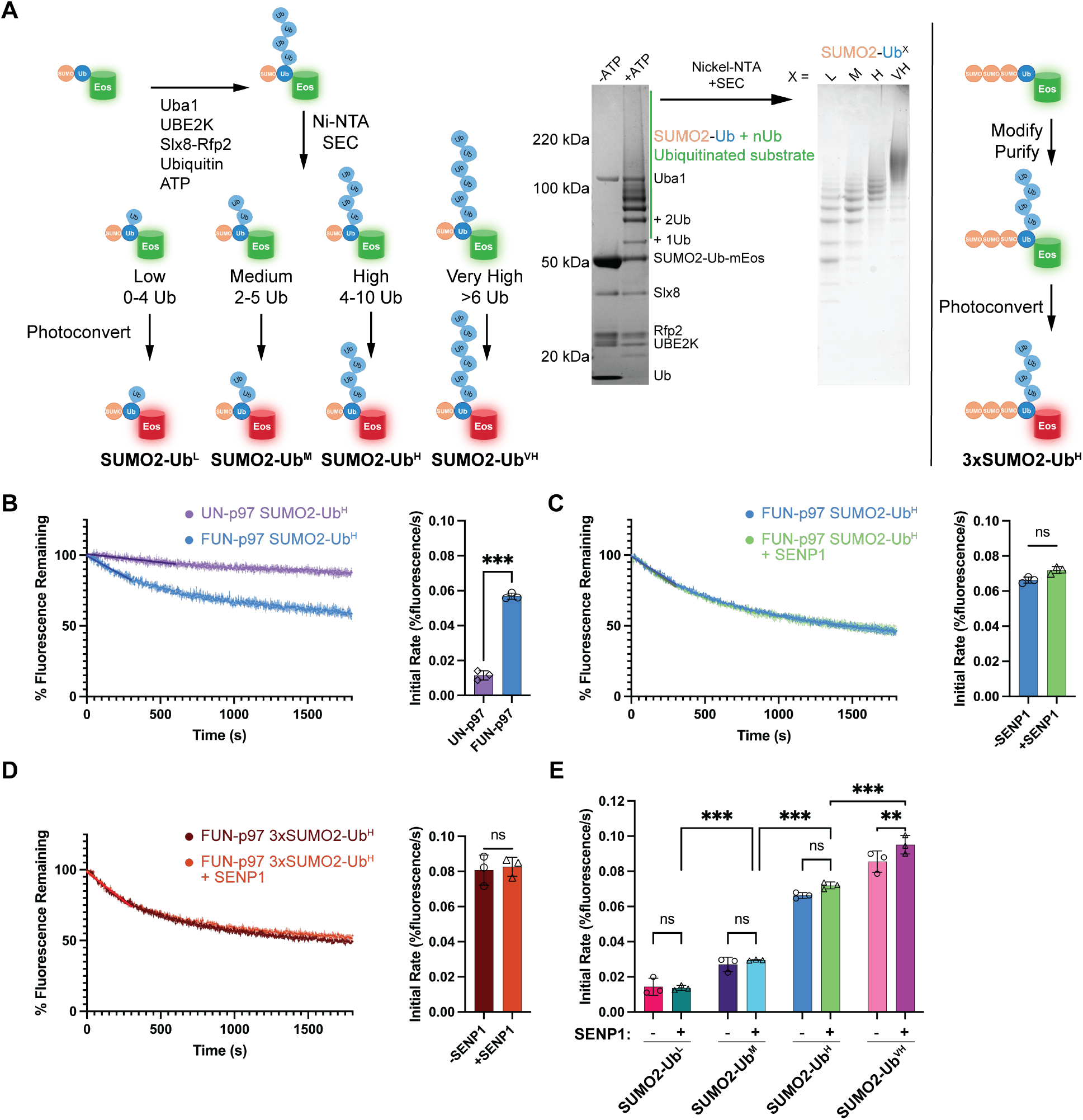
SUMO2 does not enhance unfolding by FUN-p97. **A)** *Generating SUMO-polyubiquitin conjugated substrates of varying length*. Substrates formed from a linear fusion of SUMO2, human ubiquitin, and photoconvertible fluorescent mEos were ubiquitinated with E1 *sp*Uba1, E2 *hs*UBE2K, and STUbL *sp*Slx8-Rfp2 and purified by nickel affinity (Ni-NTA) and size exclusion chromatography (SEC). Substrates were pooled by estimated length of ubiquitin chain: Low 0-4 Ub (L); Medium 2-5 Ub (M); High 4-10 Ub (H); Very High >6 Ub (VH). Left: schematic of modification, separation by size, and photocleavage. Middle: Coomassie-stained gel of modification and SYPRO-stained gel showing final pooled substrates by length. Superscript X = L, M, H, or VH denotes chain length. Right: schematic of modification of 3xSUMO2-Ub-mEos substrate. **B)** *FUN-p97 unfolds SUMO2-Ub*^*H*^ *more efficiently than UN-p97*. Left: UN-p97 and FUN-p97 unfolding SUMO2-Ub^H^ substrate with initial linear fit. Right: initial rate of unfolding as quantified from the first 600 (UN-p97) or 300 (FUN-p97) seconds of unfolding. **C)** *The presence of SUMO2 moiety in substrate does not significantly impact the rate of FUN-p97 unfolding*. Left: FUN-p97 unfolding SUMO2-Ub^H^ +/-SENP1 with initial linear fit. Right: initial rate of unfolding as quantified from the first 300 seconds of unfolding. **D)** *Additional SUMO2 moieties do not affect FUN-p97 unfolding*. Left: unfolding 3xSUMO2-Ub^H^ substrate +/-SENP1 by FUN-p97 with fit for linear region. Right: Initial unfolding rate from linear portion of unfolding (first 300 seconds). **E)** *SUMO2 does not enhance FUN-p97 at any polyubiquitin substrate length*. Initial unfolding rate from linear region of unfolding; first 600 (L, M) or 300 seconds (H, VH). P values calculated by unpaired two-tailed t-test (**B, C, D**) or ordinary two-way ANOVA with Tukey’s test (**E**); P values: 0.12 (ns), 0.033 (*), 0.002 (**), <0.001 (***).

### FAF1 enhances unfolding of polyUb-modified substrates independent of SUMO

Fission and budding yeast Ufd1 harbor a C-terminal SIM that endows UN-Cdc48 with the ability to recognize and preferentially unfold substrates modified with SUMO-polyUb hybrid chains as compared to substrates modified with polyUb-only (42). As human UFD1 lacks a SIM, we reasoned that another cofactor may assist human p97 when recognizing SUMO-polyUb hybrid chains since ubiquitin, SUMO, and SUMO-polyUb hybrid chains are conserved from yeast and human. We focused on FAF1 first as it was suggested to interact with SUMO through two proposed SIMs (44, 45). Unfolding assays with UN-p97 and FUN-p97 and the SUMO2-Ub^H^ photoconverted substrate revealed a ~4-fold increase in initial rate of unfolding for FUN-p97 relative to UN-p97 (Fig. 1B), suggesting that FAF1 includes features that increase UN-p97 activities on this SUMO-polyUb substrate. To determine if SUMO2 was responsible for enhanced activities observed for FUN-p97, SUMO2 was removed from the substrate using SENP1. Initial unfolding rates for FUN-p97 with SUMO2-Ub^H^ or 3xSUMO2-Ub^H^ substrates +/-SENP1 suggest that enhanced activities with FAF1 are not dependent on SUMO2 as all rates are comparable (Fig. 1C, 1D). While the presence of SUMO2 did not alter unfolding, given prior reports related to polyUb chain length and FAF1 (50) we next assessed if Ub chain length altered SUMO2-polyUb substrate unfolding. Initial rates of unfolding increase with Ub chain length, but no SUMO2-dependent enhancement was observed at any Ub chain length tested, as SUMO2-Ub^L^, SUMO2-Ub^M^, and SUMO2-Ub^VH^ substrates +/-SENP1 displayed comparable rates (Fig. 1E, Fig. S2B-D). In fact, for the most extensively modified Ub chain substrate (Ub^VH^), the polyUb-only substrate appears slightly better compared to the analogous SUMO2-polyUb substrate (Fig. 1E, Fig. S2D). While it is possible that SUMO2 inhibits unfolding of this substrate, we think it more likely that the simpler topology of the polyUb substrate make it a better substrate for unfolding, consistent with prior studies (41, 42). Together, these results suggest that FUN-p97 unfoldase activities are dependent on polyUb chain length but are independent of SUMO2 or polySUMO2, at least in the context of N-terminal fusions of SUMO2 or polySUMO2 relative to the Ub chain.

### FUN-p97 unfolds ubiquitinated substrates more efficiently than UN-p97 at any polyUb chain length

Given that FUN-p97 exhibited polyUb length-dependent but SUMO2-independent unfolding activities, we next focused on determinants required for polyUb-directed unfolding by FUN-p97. Some previous and several more recent studies have shown that FAF1 can enhance unfolding by UN-p97 (50–56). To directly assess polyUb-directed unfolding, we generated substrates with a single ubiquitin K48-linked chain fused to mEos via SENP1 cleavage of SUMO2 from SUMO2-Ub-mEos after ubiquitin conjugation (Fig. 2A). The resulting mixture containing SUMO2, SENP1, and polyUb-only substrates were separated by size and pooled into low, medium, high, and very high Ub chain lengths (Fig. 2A, Fig. S1E) to remove SUMO2 and SENP1. PolyUb-only photoconverted mEos substrates are hereafter referred to as Ub^X^ with the superscript denoting length (L 0-4 Ub, M 2-5 Ub, H 4-10 Ub, or VH >6 Ub). To assess FAF1 activities in the context of unfolding the Ub^H^ substrate, initial rates were compared using p97, FAF1-p97, UN-p97 and FUN-p97. For p97 or FAF1-p97, no unfolding was observed relative to a no ATP control (Fig. 2B). While FAF1 can interact with p97 (62), it does not activate p97 unfolding in the absence of UN, consistent with prior reports (46). In contrast, addition of FAF1 to UN-p97 resulted in a ~4-fold increase in the initial rate of unfolding Ub^H^ relative to UN-p97 (Fig. 2B), consistent with the fold change seen in unfolding SUMO-Ub^H^ (Fig. 1B).

**Figure 2.**
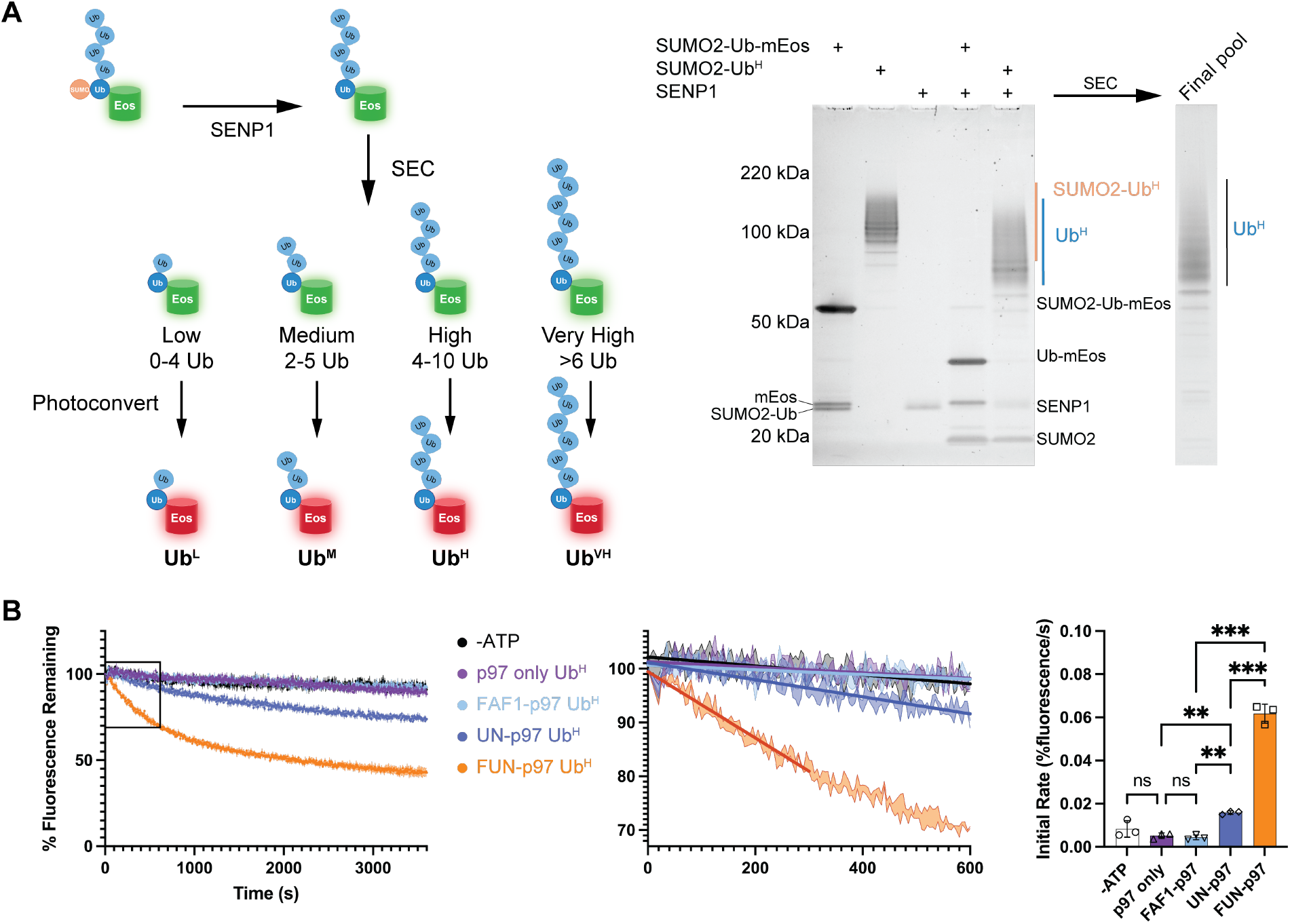
UN is required for unfolding and addition of FAF1 increases unfolding efficiency. **A)** *Removing SUMO2 to generate a polyubiquitin-only substrate*. SUMO2 is enzymatically removed using SUMO-specific peptidase 1 (SENP1) to generate Ub^X^ substrate with single ubiquitin chain. After SUMO2 removal, substrates are separated by estimated length of ubiquitin chain and photoconverted. Low 0-4 Ub (L); Medium 2-5 Ub (M); High 4-10 Ub (H); Very High >6 Ub (VH). Superscript X = L, M, H, or VH denotes chain length Left: schematic of cleavage, separation by size, and photocleavage. Right: SYPRO-stained gel showing shift in molecular weight after SENP1 cleavage for unmodified and highly modified substrate. **B)** *UN is required for unfolding; FUN-p97 unfolds substrates more efficiently*. Compared to a no ATP control, p97 only and FAF1-p97 do not unfold Ub^H^ substrate. Left: unfolding of Ub^H^ substrate by p97 only, FAF1-p97, UN-p97, and FUN-p97 with linear fit for first 600 (-ATP, p97-only, FAF1-p97, UN-p97) or 300 seconds (FUN-p97). Middle: inset showing first 600 seconds of unfolding as indicated by box on lefthand plot. Right: quantification of initial rates of unfolding from linear region. P values calculated by ordinary one-way ANOVA with Tukey’s test; P values: 0.12 (ns), 0.033 (*), 0.002 (**), <0.001 (***).

To determine relationships between ubiquitin chain length and initial unfolding rates, we assessed unfolding by FUN-p97 and UN-p97 of Ub^L^, Ub^M^, and Ub^VH^ substrates (Fig. 3A, 3B, Fig. S3). Initial rates of unfolding by FUN-p97 increase significantly with each increase in length with a 1.4-fold increase from Ub^L^ to Ub^M^, a 2.2-fold increase from Ub^M^ to Ub^H^, and a 1.7-fold increase from Ub^H^ to Ub^VH^ (Fig. 3C). In contrast, initial rates of unfolding by UN-p97 do not significantly increase between Ub^L^ and Ub^M^ or Ub^M^ and Ub^H^, but there is a significant increase of 2.6-fold when rates are compared for Ub^H^ and Ub^VH^ substrates, suggesting a threshold or minimum length of ubiquitin chain required for efficient unfolding by UN-p97 (Fig. 3C). In agreement with the observations that UBX-containing proteins can reduce the ubiquitin ‘threshold’ of UN-p97 (50), differences between unfolding activities of UN-p97 and FUN-p97 are observed for all substrates with a 2.0-fold increase observed for Ub^L^, a 2.4-fold increase for Ub^M^, a 3.7-fold increase for Ub^H^, and a 2.4-fold increase for Ub^VH^ (Fig. 3C). The contribution of FAF1 to UN-p97 unfolding appears to peak with a substrate containing 4-10 polyUb, perhaps consistent with this length representing a physiologically relevant extent of modification with polyUb (63).

**Figure 3.**
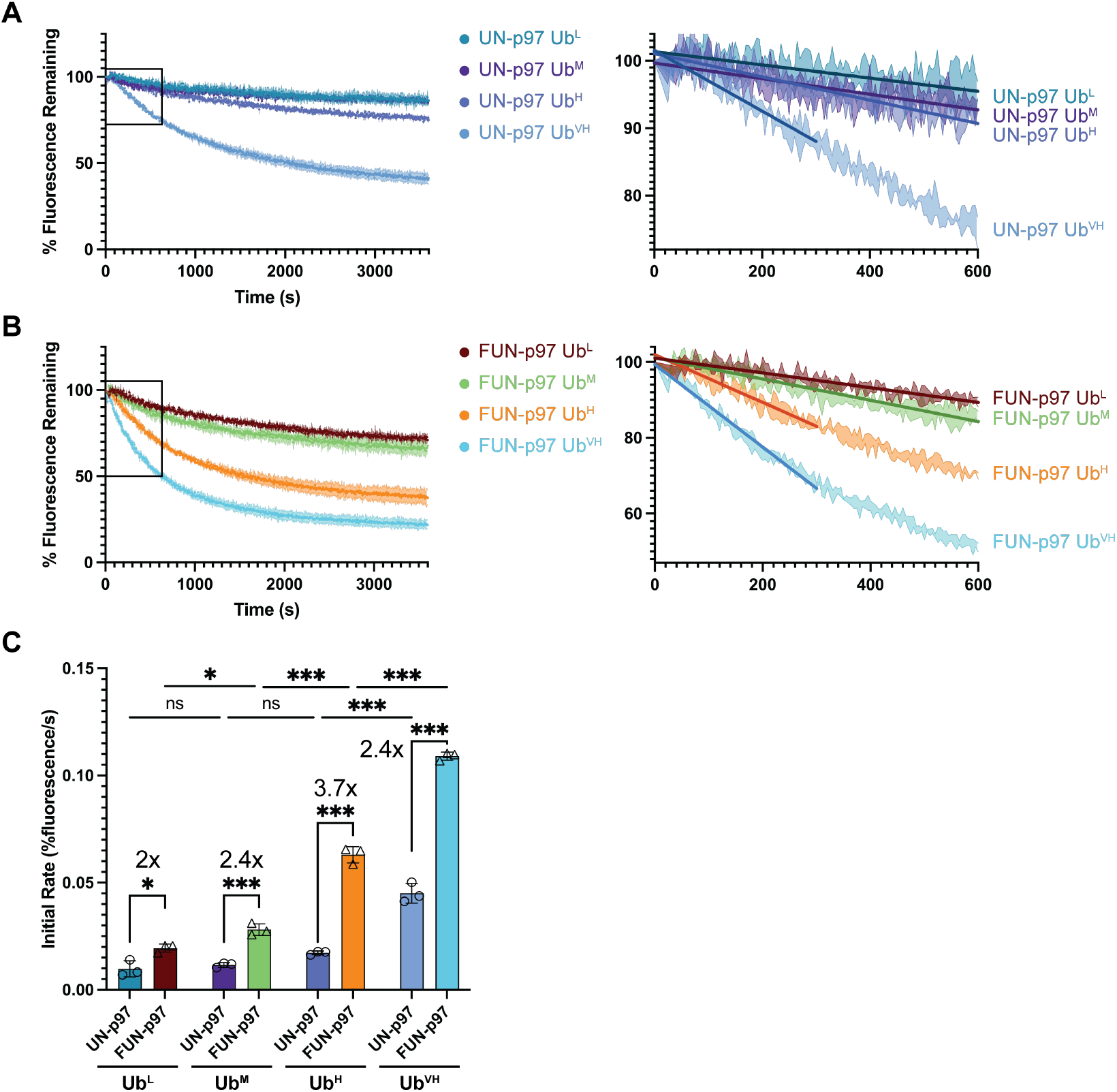
FAF1 enhances unfolding in a polyUb length-dependent manner. **A)** *UN-p97 requires longer polyUb chains for efficient unfolding*. Left: UN-p97 unfolding all substrate lengths. Right: first 600 seconds of UN-p97 unfolding as indicated by box in lefthand plot with linear fits for first 600 (L, M, H) or 300 seconds (VH). **B)** *FUN-p97 unfolds longer ubiquitin chains faster*. Left: FUN-p97 unfolding all substrate lengths. Right: first 600 seconds of FUN-p97 unfolding as indicated by box in lefthand plot with linear fits for first 600 (L, M) or 300 seconds (H, VH). **C)** *Rate of unfolding increases in the presence of FAF1*. Quantification of initial rate of unfolding determined from slopes of first 600 (L, M, UN-p97 H) or 300 seconds (FUN-p97 H, VH) of unfolding. Fold-change between UN-p97 and FUN-p97 at each length is indicated. P values calculated by ordinary two-way ANOVA with Tukey’s test; P values: 0.12 (ns), 0.033 (*), 0.002 (**), <0.001 (***).

### CryoEM reconstructions of FUN-p97 bound to ubiquitinated substrate

To gain structural insight into FAF1 and its activities in unfolding by FUN-p97, we determined single particle cryoEM reconstructions for FUN-p97 in the presence of ATP and the Ub^H^-mEos unconverted green substrate. To interpret these reconstructions, we used recently determined atomic models of FUN-p97 with two truncated FAF1 constructs containing the UBX domain and partial helical domains in the presence (PDB 11TA) and absence (PDB 11VE) of ubiquitinated substrate (56) in conjunction with AlphaFold3 (64) predictions for a truncated FAF1 containing the UBX, helical, and UAS domains binding to the UFD1 UT3 domain to fit our reconstructions (Fig. 4A). We rigid-body fit PDBs 11TA and 11VE in our densities, docked the truncated FAF1-UFD1 AlphaFold on the UBX-bound p97 N-domain, and colored to match. Our reconstructions were determined with nominal resolutions of 4.16-4.74 Å, but due to heterogeneity and disorder and given recent high-resolution structures of UN-p97 with truncated FAF1 (56), we present depictions of our reconstructions as low pass filtered images at 6 Å, unless otherwise noted, to enable comparisons and to highlight similarities and differences between observed conformations.

**Figure 4.**
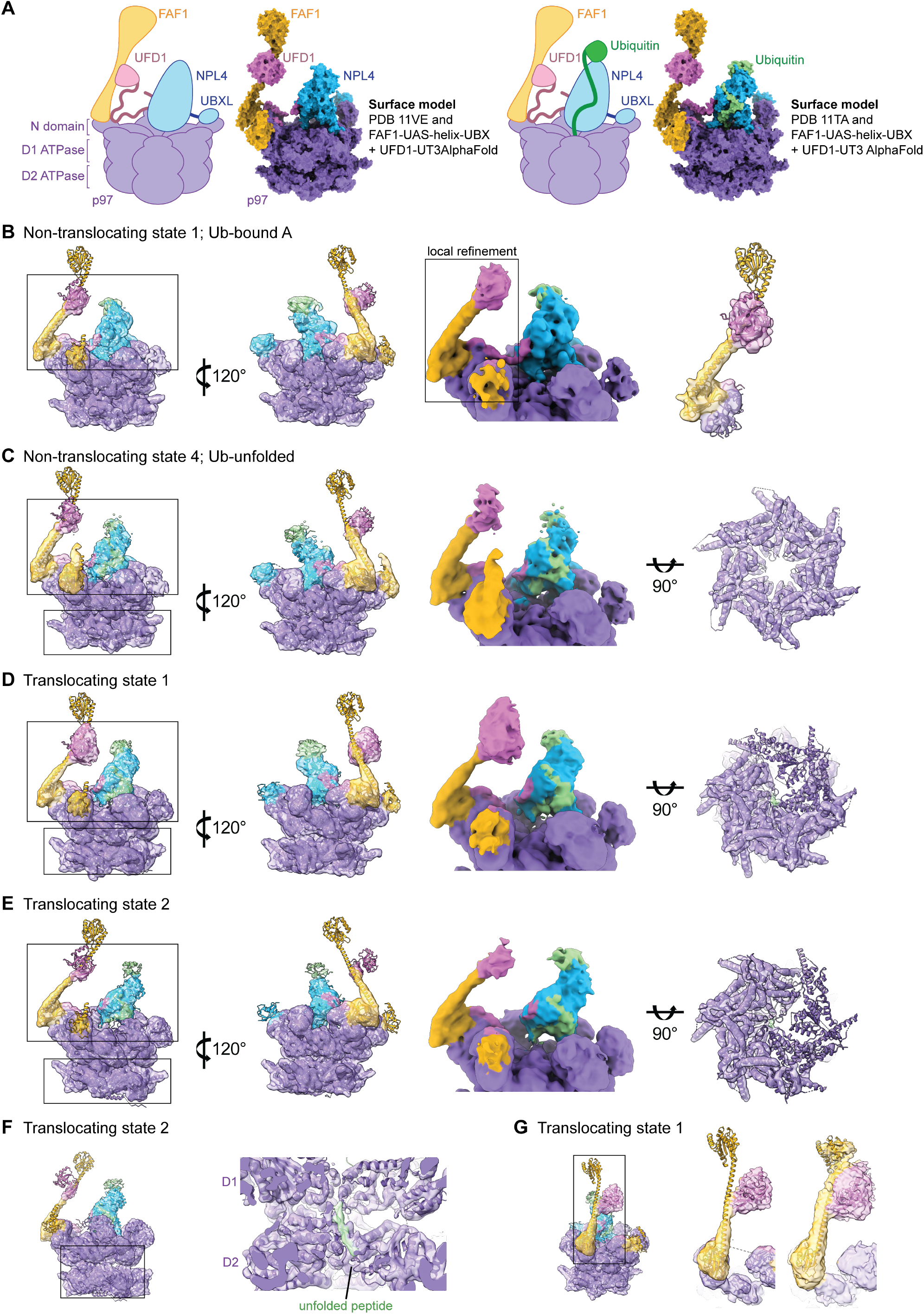
Cryo-EM reconstructions for FUN-p97 actively unfolding a ubiquitinated substrate. **A)** *Model of FUN-p97 in the absence and presence of ubiquitinated substrate*. Left: cartoon depiction of FUN-p97 complex and surface model of PDB 11VE with AlphaFold prediction of FAF1-UAS-helix-UBX + UFD1-UT3 docked onto UBX-bound N-domain of p97. Right: cartoon depiction of FUN-p97 complex bound to ubiquitin substrate and surface model of PDB 11TA with AlphaFold prediction of FAF1-UAS-helix-UBX + UFD1-UT3 docked onto UBX-bound N-domain of p97. Throughout the figure cartoons, ribbon diagrams, and density maps are colored as follow: FAF1 (yellow), UFD1 (pink), NPL4 (blue), p97 (purple), ubiquitin (green). **B)** *Non-translocating state 1 of FUN-p97 prior to ubiquitin-unfolding low pass filtered to 6 Å; ‘Ub-bound A’*. Left: AlphaFold model FAF1-UAS-helix-UBX, PDB 11VE (p97, UFD1, NPL4, FAF1 UBX), and PDB 1UBQ (Ub) rigid body fit into two views of electron density map. Middle: Close-up view of interactions between NPL4, UFD1, FAF1, and p97. Right inset: focused refinement of p97 N-domain and FAF1 density fit with AlphaFold FAF1-UAS-helix-UBX + UFD1-UT3. **C)** *Non-translocating state 4 of substrate-bound FUN-p97 with ubiquitin unfolded in NPL4 channel low pass filtered to 6 Å; ‘Ub-unfolded’*. Left: AlphaFold model FAF1-UAS-helix-UBX, PDB 11TA (p97, UFD1, NPL4, FAF1 UBX, unfolded Ub), and PDB 1UBQ (folded Ub) rigid body fit into two views of electron density map. Middle: Close-up view of interactions between NPL4, UFD1, FAF1, ubiquitin, and p97. Right: bottom of p97 D2 ring from refinement prior to low pass filtering. **D)** *Translocating class of substrate-bound FUN-p97 with ubiquitin unfolded in NPL4 channel and density of unfolded ubiquitin in p97 pore low pass filtered to 6 Å; ‘Translocating state 1’*. Left: AlphaFold model FAF1-UAS-helix-UBX, PDB 7LN1 (p97), PDB 11TA (UFD1, NPL4, FAF1 UBX, unfolded Ub), and PDB 1UBQ (folded Ub) rigid body fit into two views of electron density map. Middle: Close-up view of interactions between NPL4, UFD1, FAF1, Ub, and p97. Right: bottom of p97 D2 ring from refinement prior to low pass filtering. **E)** *Translocating class of substrate-bound FUN-p97 with ubiquitin unfolded in NPL4 channel and density of unfolded ubiquitin in p97 pore with NPL4 exhibiting a different conformation low pass filtered to 6 Å; ‘Translocating state 2’*. Left: AlphaFold model FAF1-UAS-helix-UBX + UFD1-UT3, PDB 7LN1 (p97), PDB 11TA (UFD1, NPL4, FAF1 UBX, unfolded Ub), and PDB 1UBQ (folded Ub) rigid body fit into two views of electron density map. Middle: Close-up view of interactions between NPL4, UFD1, FAF1, ubiquitin, and p97. Right: bottom of p97 D2 ring from refinement prior to low pass filtering. **F)** Left: Density map of translocating state 2 fit with AlphaFold model FAF1-UAS-helix-UBX, PDB 7LN1 (p97), PDB 11TA (UFD1, NPL4, FAF1 UBX, unfolded Ub), and PDB 1UBQ (folded Ub). Right: inset showing unfolded substrate peptide in p97 pore in density map prior to low pass filtering. **G)** Left: Density map of translocating state 1 fit with AlphaFold model FAF1-UAS-helix-UBX + UFD1-Ut3, PDB 7LN1 (p97), PDB 11TA (UFD1, NPL4, FAF1 UBX, unfolded ubiquitin), and PDB 1UBQ (folded Ub). Middle: Close-up view of FAF1 and UFD1 interactions. Left: Lower contour view, fit with only AlphaFold model FAF1-UAS-helix-UBX + UFD1-UT3.

We incubated FUN-p97 containing full-length FAF1, Ub^H^-mEos unconverted green substrate, and ATP• Mg for 5-10 minutes before application to grids and vitrification in an attempt to capture complexes during unfolding (~50% of substrate unfolded at this time range). Three separate datasets were combined to obtain ~2 million particles. After discarding junk and particles containing just p97 or p97 dodecamer, ~900k particles were obtained with densities consistent with UN and FAF1 on top of p97 with p97 ‘N-domain-up’ conformation, indicative of ATP-bound D1 protomers and an increase in cofactor affinity (29, 65, 66).

Particles were next segregated into non-translocating (NT) (67.7%) and translocating (32.3%) classes using hetero refinement prior to subsequent rounds of 3D classification. The NT class was further separated into 10 classes using 3D classification without image alignment which show varying positions and quality of densities for NPL4 and FAF1 (Fig. S4). Four NT classes contained stronger densities for both NPL4 and at least one FAF1, which we assigned as NT state 1 ‘Ub-bound A’ (Fig. 4B), NT state 2 ‘Ub-bound B’ (Fig. S5A), NT state 3 ‘Unbound’ (Fig. S5B), and NT state 4 ‘Ub-unfolded’ (Fig. 4C). All NT classes exhibit six-fold symmetry with respect to the planar conformation of p97 D1 and D2 rings, suggesting that these classes are not actively unfolding substrate within the pore (Table 1, Fig. 4C, right inset). Consistent with this, NT classes Ub-bound A, Ub-bound B, and Unbound reveal densities for the NPL4 loop (residues 425-439) which obstructs the central pore of p97 (42, 56, 66). Densities atop NPL4 are consistent with ubiquitin within the Ub-bound and Ub-unfolded classes as based on prior structures (39, 42, 56, 67) and the position of NPL4 varies by approximately 25 degrees between the Ub-bound and Unbound states relative to the central pore of p97 (Fig. S5C) in agreement with the ‘seesaw’ conformation of NPL4 previously reported (67). Two FAF1 molecules appear bound to p97, although one exhibits stronger densities consistent with p97-bound FAF1 UBX domain and an extended FAF1 helix while the other exhibits weaker densities for a second UBX domain on an adjacent N domain in some states, including that of the Ub-unfolded reconstruction (Fig. 4C). Previous work (56) combined with rigid body fitting AlphaFold3 predictions for FAF1-UBX-helix-UAS bound to UFD1 UT3 suggests that UFD1 interacts with the helical domain of FAF1 in multiple classes (Fig. 4, Fig. S5). Focused refinement of Ub-bound A on regions including the p97 N-domain, FAF1, and UFD1 did not reveal additional densities for FAF1 UAS domain or other ubiquitin binding domains (Fig. 4B, right inset).

**Table 1.** Positions of FAF1/UFD1, NPL4, and p97 in cryo-EM reconstructions.

| Reconstruction | FUN-p97 NT state 1<br>Ub-bound A | FUN-p97 NT state 2<br>Ub-bound B | FUN-p97 NT state 3<br>Unbound | FUN-p97 NT state 4<br>Ub-Unfolded | FUN-p97 Translocating state 1 | FUN-p97 Translocating state 2 |
| --- | --- | --- | --- | --- | --- | --- |
| **FAF1/UFD1** | Forward | Back | Middle | Back | Forward | Forward |
| **NPL4 Tower** | Back | Back | Forward 2 | Back | Forward 1 | Middle |
| **NPL4 loop** | Closed | Closed | Closed | Open | Open | Open |
| **D1** | Symmetric | Symmetric | Symmetric | Symmetric | Staircase | Staircase |
| **D2** | Planar | Planar | Planar | Planar | Staircase | Staircase |

FAF1 adopts at least two conformations in NT classes, hinging at the UBX domain, with the helix moving approximately 10 degrees relative to p97 between our Ub-unfolded and Ub-bound A classes (Fig. S5D) and 9 degrees relative to p97 compared to the FAF1 position in previous FUN-p97 reconstructions (Fig. S6A) (56), perhaps to position the UT3 of UFD1 for substrate engagement and unfolding initiation.

In addition to the Ub-bound A, Ub-bound B, and Ub-unfolded states which are comparable to previously observed non-translocating FUN-p97 ubiquitin-bound initiation states (56), we isolated FUN-p97 particles and reconstructions representing translocating states as evidenced by an open central pore, non-planar D1 and D2 rings with protomers in the D2 ring adopting a spiraling conformation, and contacts with substrate within the central pore (Table 1) (39, 68). To interpret reconstructions for translocating p97, PDB 7LN1 was used for rigid body fitting with PDB 11TA used to dock components of FUN into reconstructions. Substrate peptide engaged within the D2 ring was observed in reconstructions of translocating classes (Fig. 4D-F insets). A consensus reconstruction for the translocating class exhibits weaker densities for NPL4 and FAF1 (Fig. S4), suggesting multiple conformations for cofactors within the translocating classes. Particles were separated into 6 classes using 3D classification without image alignment to reveal two classes with stronger densities for NPL4, FAF1, UFD1, and ubiquitin. We designate these translocating classes as ‘Translocating state 1’ (T1) (Fig. 4D) and ‘Translocating state 2’ (T2) (Fig. 4E). The T1 and T2 reconstructions are different with respect to the location of substrate peptide bound to D2 and p97 protomers with reduced densities for D2 rings that is thought to represent the protomer undergoing active (or after) ATP hydrolysis (see Fig. 4D, E right insets) (39, 69). NPL4 in the T1 class adopts a conformation distinct from previously reported FUN-p97 NPL4 states (56), with NPL4 of T1 approximately aligned with the central pore of p97 and an NT initiation complex leaning 13 degrees behind the central pore relative to T1 and an NT pre-initiation complex leaning 17 degrees in front of the central pore of p97 relative to T1 (Fig. S6B). The NPL4 position of T1 is also distinct from NPL4 positions in our observed NT states (Table 1). T2 represents a conformation of NPL4 that leans away from FAF1 and UFD1 by approximately 16 degrees relative to T1 (Fig. S5E) and leans 29 or 2 degrees away from the central pore of p97 relative to NT initiation and pre-initiation complexes, respectively (Fig. S6C). Both positions are similar to those previously seen in the ‘see-sawing’ of NPL4 in UN-p97 complexes (67). T1 and T2 both have densities for FAF1 in a position similar to the FAF1 position seen in Ub-bound A (Fig. S5D, Fig. S5F). This position is distinct from the FAF1 position in Ub-unfolded (Fig. S5F), with FAF1 leaning 9 degrees relative to the position of FAF1 in Ub-unfolded with a fulcrum at the UBX domain, suggesting that FAF1 adopts different conformations throughout substrate binding, unfolding initiation, and active unfolding, as seen in our translocating and non-translocating reconstructions.

Reconstructions of T1 reveal the strongest density for FAF1 and UFD1. In low pass filtered reconstructions densities adjacent to the FAF1 helix appear larger than UFD1’s UT3 (Fig. 4D) suggesting that ubiquitin may be bound to UFD1 as previously observed (56). In higher resolution reconstructions, densities were observed for a portion of the FAF1 helix and UFD1 UT3, and at lower contour additional densities on the side of the helix and beyond UFD1 become clear, possibly representing the FAF1 UAS domain (Fig. 4G, Fig. S6D, E) which we were otherwise unable to resolve.

### The helix-UBX motif of FAF1 is sufficient to promote unfolding

Reconstructions presented here alongside AlphaFold3 models (64) and recently reported structures (56) are consistent with interactions between the FAF1 UBX domain and p97, the FAF1 helix and UFD1, and include additional densities which may represent other functional domains of FAF1. Recent studies have used analogous assays to dissect the role of FAF1 domains (51–53, 56), but these studies were conducted using substrates which were extensively modified with >6 Ub and/or Ub-Ub-mEos substrates which generate branched chains when modified which are more complex to unfold. As such, we chose to test analogous FAF1 truncations to determine if any domains impact unfolding under limiting polyUb conditions by comparing Ub^VH^, Ub^H^, Ub^M^, and Ub^L^ unfolding rates for different FAF1 constructs.

The FAF1 helix interacts with UFD1 as predicted by AlphaFold and later confirmed by recent studies (51– 56). Further, the UBA domain of FAF1 was implicated in ubiquitin interaction (49, 70). To determine if these domains become more important when polyUb is limiting in our Ub^L^, Ub^M^, and Ub^H^ substrates, FAF1 constructs were generated to remove the ubiquitin binding domain (ΔUBA, residues 48-650) or containing only the helical and p97-binding UBX domain (helix-UBX, residues 471-650) (Fig. 5A). Unfolding assays reveal that FAF1-dependent activities are maintained in constructs lacking the UBA domain (ΔUBA) or lacking the UBA, ubiquitin-like (UBL) 1 and 2, and UAS domains (helix-UBX) independent of polyUb chain length as initial rates for each of these complexes is higher than UN-p97 and similar to those observed for FUN-p97 at each length of polyUb chain (Fig. 5A, 5B; Fig. S7). Given the helix-UBX is sufficient to increase initial unfolding rates, the UBL and fatty-acid-binding (UAS) (71) domains, in addition to the UBA domain, appear to be dispensable for increased unfolding. Together with our structural data and prior results (51–56) our biochemical data are consistent with a model suggesting that rather than FAF1 binding to ubiquitin, increased activity is due to UFD1 interactions with the helical domain of FAF1 which in turn stabilizes cofactor and substrate association with the p97 complex (Fig. 5C).

**Figure 5.**
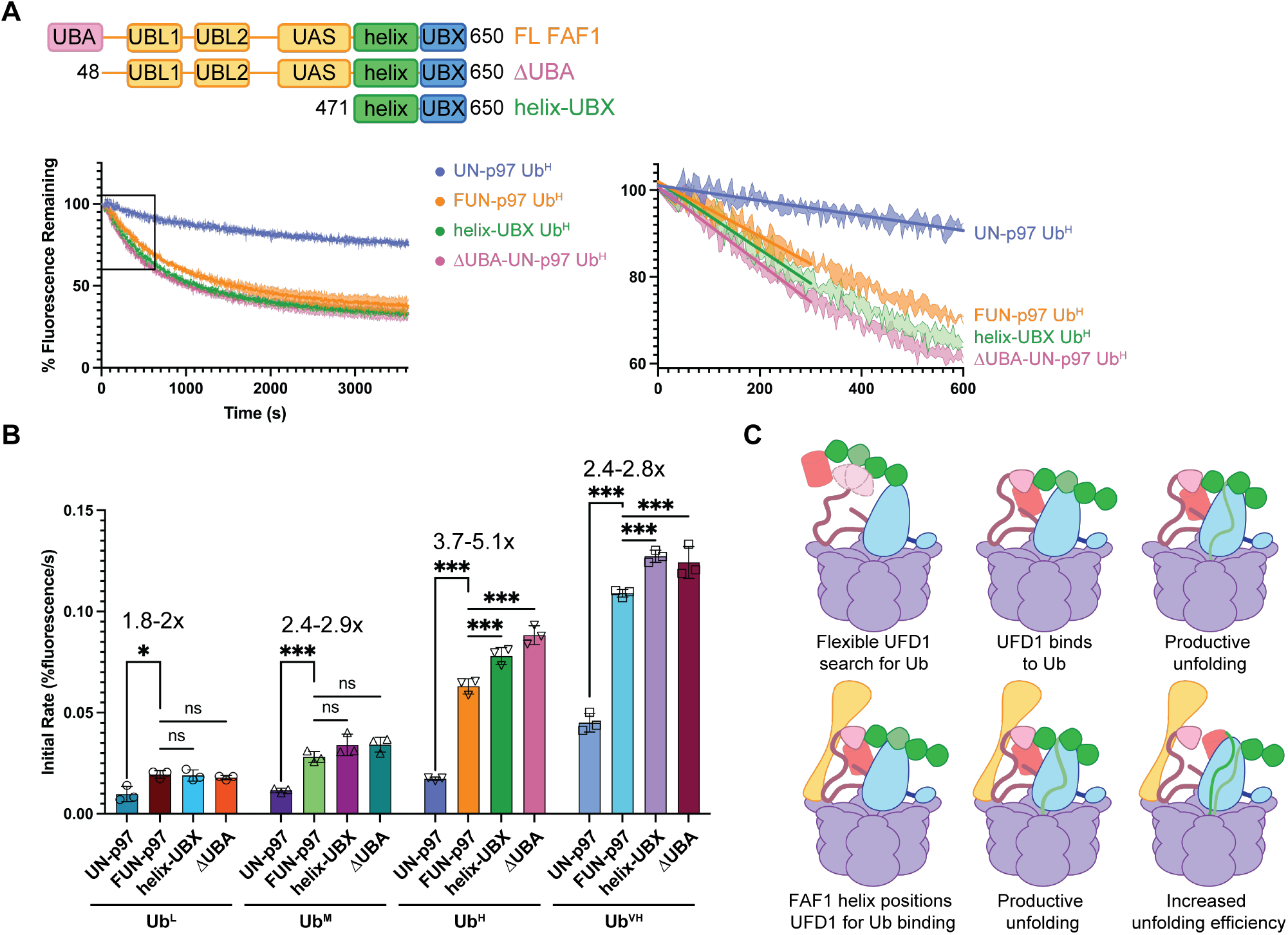
FAF1 promotes unfolding via helix-UBX motif. **A)** *Deletion of ubiquitin-associated (UBA) domain (ΔUBA) has no effect on FUN-p97 unfolding; the helical and p97-binding (UBX) domains of FAF1 (helix-UBX) appear to be sufficient to induce the increase in unfolding relative to UN-p97*. Top left: FAF1 domains and constructs with domains deleted. Bottom left: unfolding Ub^H^ substrate by UN-p97, FL FUN-p97, or mutant FUN-p97 with helix-UBX or *Δ*UBA FAF1 constructs. Right: first 600 seconds of unfolding as indicated by box in lefthand plot with linear fits. **B)** *Length of substrate does not affect role of helix-UBX*. Quantification of initial rate of unfolding from first 600 (L, M) or 300 seconds (H, VH). Range of fold change from UN-p97 to FUN-p97/mutants indicated. P values calculated by ordinary two-way ANOVA with Tukey’s test; P values: 0.12 (ns), 0.033 (*), 0.002 (**), <0.001 (***). **C)** Model for UN-p97 unfolding (top) vs FUN-p97 unfolding (bottom) Colored as follows: FAF1 (yellow), UFD1 (pink), NPL4 (blue), p97 (purple), ubiquitin (green), initiator ubiquitin (light green), mEos substrate (red).

## DISCUSSION

We set out to determine if FAF1 contributes to recognition of substrates modified by SUMO-polyUb hybrid chains. However, we observed no effect of SUMO2 on UN-p97 or FUN-p97 unfolding in the context of SUMO2-polyUb or polySUMO2-polyUb modified substrates. While it remains possible that the topology of our substrates does not represent a configuration that leads to productive SUMO2-polyUb-directed activity, it is also possible that any SUMO interaction via FAF1 could result in binding that does not enhance unfolding rates. Our study did not include SUMO1-containing substrates, so it is also possible that FAF1 recognition is specific to the chain-capping SUMO variant (44, 72, 73). While SUMO2/3 are more commonly associated with polyUb, a STUbL specific for SUMO1-capped SUMO2/3 chains was reported (74), so it remains possible that a combination of SUMO variants may be required for chain recognition. While the aforementioned points leave open the possibility for FAF1-dependent SUMO recognition for different chain topologies, other p97 cofactors may serve as human SUMO-polyUb readers.

Our biochemical data show that FAF1 increases initial rates of unfolding relative to UN-p97 at all lengths of ubiquitin chain tested, consistent with previous reports that FAF1 is able to reduce the polyUb threshold (50) to productively unfold substrates with shorter polyUb chains that UN-p97 does not efficiently unfold. Interestingly, the effects of FAF1 appear greatest for polyUb chains containing between 4 and 10 ubiquitin moieties, which is more representative of the majority of ubiquitin chains in cells (63, 75).

Our structures and biochemistry are fully consistent with recent studies that suggest that the helical domain of FAF1 binds to and potentially stabilizes the UT3 domain of UFD1 (51–56). Our work extends prior reports in that we were able to isolate FUN-p97 complexes in translocating states in addition to non-translocating states, revealing the respective positions of FAF1, UFD1, and NPL4 during active translocation. Notably, in the presence of FAF1, UFD1 appears to be anchored and positioned closer to NPL4 and the site of ubiquitin unfolding initiation throughout substrate engagement, unfolding initiation, and active translocation, compared to UN-p97 structures where UFD1 is not resolved and presumably labile (39, 40, 42, 67, 69). The positions of FAF1, UFD1, and NPL4 shift relative to one another throughout translocation, with UFD1/NPL4 core appearing to move away from p97 by 6.4 and 8.7 Å during translocation, in T1 and T2 respectively, as measured by displacement at NPL4 glutamine 390 (Fig. S6F). This displacement captures movement/displacement due to spiraling of the D1 ring in addition to upward motion of the NPL4 tower, as well as rotation of the tower relative to D1. Densities for NPL4 also become more disordered during translocation, further suggesting that the role of FAF1 may be to hold onto the UFD1 UT3 domain during these transitions to ensure that UN is able to dock back onto the p97 surface after each step in translocation. While the UBA, UAS, or UBL domains are unresolved for FAF1 in our reconstructions, it remains possible that they interact with ubiquitin or other polyUb topologies (or SUMO-polyUb topologies) to further enhance activities not tested in this study. As such, functions for FAF1 and other p97 adaptors will require further efforts to determine substrate specificities and Ub/Ubl topologies that more accurately reflect their physiologic substrates and binding partners.

## METHODS

### Protein Expression and Purification

SENP1Δ419 (76), *sp*Uba1 (77), *hs*UBE2K (78), and *sp*Slx8-Rfp2 (42) were expressed and purified as previously described. Human ubiquitin was cloned into pET3C using Gibson Assemby (NEB) and expressed and purified as previously described (79). The Gibson Assembly kit (NEB) was used to generate linear fusions of SUMO2, ubiquitin, and the mEos3.2 gene (Addgene) inserted into pET28-NTEV to generate SUMO2-Ub-mEos. The 3xSUMO2-Ub-mEos construct was purchased from GenScript in pET28a(+)-TEV. Fluorescent fusion proteins were expressed and purified as previously described (42). Codon optimized UFD1 was inserted into the pTrx28 vector (80), codon optimized NPL4 was inserted into MCS2 of pRSF-Duet1, codon optimized p97 and FAF1 were inserted into pET28-NTEV, and plasmids containing FAF1 mutants; helix-UBX aa471-650, ΔUBA aa48-650 were ordered from GenScript in pET28a(+) vector with a TEV-cleavable Trx included in the sequence for insertion. Plasmids were transformed into *E. coli* BL21-CodonPlus (DE3)-RIL cells for expression. Cells were grown at 37°C in Super Broth medium (Teknova) until OD600 reached 1.0. To induce expression, IPTG was added to 0.3 mM and cells were moved to 18°C overnight. Cells were harvested by centrifugation and resuspended in lysis buffer containing 1µg/L DNAse, 1 mg/L lysozyme, 20 mM HEPES pH 7.5, 20 mM imidazole 0.1% IGEPAL CA-630, 0.1 mM TCEP, 10 mM MgCl2, and 20% sucrose with 150 mM NaCl for UFD1/NPL4/p97, and 500 mM KCl for FAF1. UFD1 and NPL4 cells were combined at the lysis step for purification. Lysate was sonicated and centrifuged for 25 minutes at 45,000x g. Supernatant was applied to Ni-NTA agarose beads (ThermoFisher) after equilibration with a wash buffer containing 20 mM HEPES pH 7.5, 20 mM imidazole, and salt as listed previously. Beads were washed with 5 column volumes of wash buffer and proteins eluted with 3 column volumes elution buffer (300 mM imidazole). Eluted proteins were dialyzed in the presence of TEV protease and separated using HiLoad 26/60 Superdex 200 column (Cytiva) or HiLoad 26/60 Superdex 75 column (Cytiva) in 20 mM HEPES pH 7.5, 0.1 mM TCEP and 150 mM NaCl for UFD1/NPL4/p97 or 500 mM KCl for FAF1 constructs. Fractions containing proteins of interest were pooled and flash frozen in liquid nitrogen and stored at −80C.

### mEos Substrate Ubiquitination

Conjugation reactions for indicated base substrate included 0.5 µM *sp*Uba1, 2 µM *hs*UBE2K, 2 µM *sp*Slx8-Rfp2, 5 µM substrate, and 50 µM human ubiquitin incubated in 50 mM HEPES pH 7.5, 150 mM NaCl, 5 mM MgCl_2_, 0.1 mM TCEP, and 5 mM ATP for 2 hours at 37°C. Imidazole was added to a final concentration of 20 mM prior to application to Ni-NTA beads. The beads were washed with 5 column volumes of wash buffer (20 mM HEPES pH 7.5, 150 mM NaCl, 20 mM imidazole), and substrate was eluted with 3 column volumes of elution buffer (20 mM HEPES pH 7.5, 150 mM NaCl, 300 mM imidazole). Eluates were diluted to a final concentration of 100 mM NaCl and applied to MonoQ 5-50 GL (Cytiva) equilibrated with 100 mM NaCl, 20 mM HEPES pH 7.5, 0.1 mM TCEP and eluted with a gradient to 50% of 1 M NaCl, 20 mM HEPES pH 7.5, 0.1 mM TCEP. Fractions containing the protein of interest were pooled and flash frozen in liquid nitrogen and stored at −80C. To generate Ub-only substrates 5 equivalents of SENP1Δ419 was added immediately following ubiquitin conjugation and incubated at 4°C for 18 hours. Protein was further fractionated by size-exclusion chromatography (S200 Increase) equilibrated with 150 mM NaCl, 20 mM HEPES pH 7.5, 0.1 mM TCEP to separate proteins by polyubiquitin length. Fractions containing proteins of interest were pooled and flash frozen in liquid nitrogen and stored at −80C.

### Substrate Photoconversion

Polyubiquitinated mEos substrates were irradiated at 365 nm on ice with the ENF-280C transilluminator (Spectroline) (81) after ubiquitination and separation by polyubiquitin length. Substrates were irradiated until half of the proteins were cleaved, as measured by decrease in molarity of green substrate via 506 nm absorbance. Unless specified, substrate concentrations refer to photocleaved concentration measured by 571 nm absorbance.

### Unfoldase Assay

400 nM hexameric p97, 400 nM UFD1/NPL4, 400 nM FAF1 FL or mutant (as specified), and 40 nM photocleaved substrate as specified was incubated in 5 mM KCl, 50 mM Tris pH 7.5 at 4C, 20 mM MgCl_2_, 1 mM EDTA, 30 mM creatine phosphate, 3 U/mL phosphokinase, and 3 U/mL pyrophosphatase at 37°C for 10 minutes alone or in the presence of 10 µM SENP1 where indicated. Reactions were initiated by the addition of 5 mM ATP and monitored using a SpectraMax M5*e* and SoftMax Pro 5 software (Molecular Devices).

Fluorescence at an excitation wavelength of 540 nm and an emission wavelength of 570 nm was measured every 5 seconds for 1 hour or 30 minutes (+/-SENP1 assays) at 37°C. The ‘percent substrate remaining’ was calculated by normalizing the data and determining 100% as a t0 value calculated from linear regression of the first 120s of unfolding. Initial rates were determined using a linear fit to the initial linear portion of the data; 600 seconds for L, M, and UN H substrates or 300 seconds for VH and FUN H substrates using Prism 11 (GraphPad).

### Cryo-Electron Microscopy Sample Preparation and Data Collection

FUN-p97 and substrate were incubated together (2 µM FAF1, 2 µM UFD1/NPL4, 2 µM hexameric p97, 2 µM Ub^H^ non-converted substrate) in 5 mM KCl, 50 mM Tris pH 7.5 at 4C, 20 mM MgCl_2_, 1 mM EDTA, 30 mM creatin phosphate, 3 U/mL pyrophosphatase, 3 U/mL phosphokinase at 37°C prior to addition of 5 mM ATP and incubating at 37°C for 4 min 45 seconds (dataset 1), 9 min 15 seconds (dataset 2), or 8 min 45 seconds (dataset 3). CHAPSO was added to a final concentration of 0.05% immediately prior to vitrification. 4 µL of sample was applied to a glow-discharged UltrAufoil, 1.2/1.3, 300 mesh grid (Quantifoil) in 100% humidity at 22°C, blotted for 4s after 8s wait, and plunged into liquid ethane using the FEI Vitrobot Mark IV (ThermoFisher). Data collection was carried out using a Titan Krios 300KV (FEI) instrument equipped with a K3 Summit direct detector (Gatan). 6,566, 5,653, and 7,771 movies (40 frames/movie, 4 s exposure time) were collected in three sessions in super-resolution mode with a defocus range from −0.8 to −2.3 μm at a dose rate of ~ 20 e^−^/px/sec and a total dose of 70.7 e^−^/Å^2^/movie. The calibrated pixel size was 1.064 Å/px.

### Image Processing

A summary of image processing steps is shown in Fig. S4. A total of 19,990 movies were imported into cryoSPARC Live for motion correction and CTF estimation, subsequent processing steps were all completed in cryoSPARC 4.7 (82). Movies with CTF estimate between 2 Å and 6 Å and with ice thickness between 0.958 and 1.08 were discarded for a total of 16,572 remaining movies. A total of 2,075,500 particles were selected via unsupervised blob picking and subjected to 2D classification and to remove junk particles. The remaining 1,958,674 particles were subject to heterogenous refinement using ab initio reconstructions as initial volumes. Particles containing junk or only p97 were removed and the remaining 953,556 were re-extracted from micrographs with a box size of 512 pixels for a stack of 898,460 particles containing densities for p97, UN, and FAF1. After subsequent heterogeneous refinement, the particle stack was segregated into translocating (290,178 particles) and non-translocating (303,543 particles) particle stacks. Classes were subjected to further 3D classification without image alignment and subsequent non-uniform refinement. Four non-translocating classes revealed strong densities for UN and FAF1 and we have designated them NT states 1-4, ‘Ub-bound A’, ‘Ub-bound B’, ‘Ub-unfolded’, and ‘Unbound’. Ub-bound A was further refined using local refinement with a mask around densities for FAF1, UFD1, and a portion of p97 N-domain. Two translocating classes revealed strong densities for UN and FAF1 and we have designated them ‘Translocating state 1’ and ‘Translocating state 2’. The AlphaFold3-predicted structural model of truncated FAF1 containing UBX, helical, and UAS domains binding to UT3 domain of UFD1 was docked into densities along with PDBs 11TA, 11VE, and 1UBQ using ChimeraX (83). Local resolutions of reconstructions were determined using PHENIX v1.19.2-4158 (84).

## DATA AVAILABILITY

EM reconstructions are deposited in the Electron Microscopy Data Bank (EMDB) with the following accessing codes: FUN-p97 NT state 1 Ub-bound A, EMD-78389; FUN-p97 NT state 2 Ub-bound B, EMD-78396; FUN-p97 NT state 3 Unbound, EMD-78395; FUN-p97 NT state 4 Ub-unfolded EMD-78390; FUN-p97 Translocating state 1 EMD-78397; FUN-p97 Translocating state 2 EMD-78391, and will be released upon review by RCSB staff.

## ACKNOWLEDGEMENTS

We thank members of the Lima lab for advice. We thank Alessandro Cirulli for assistance in purifying FAF1 ΔUBA and helix-UBX constructs. We thank Jason De La Cruz and Sagnik Sen for input at early stages of cryo-EM data collection and processing. The content is solely the responsibility of the authors and does not represent the official views of the National Institutes of Health. C.D.L. is an investigator of the Howard Hughes Medical Institute.

## AUTHOR CONTRIBUTIONS

**Abigail A. Lemmon:** Conceptualization (co-lead); Investigation (lead); Writing – original draft (lead).

**Christopher D. Lima:** Conceptualization (co-lead); Investigation (advised); Writing – editing.

