## Supporting Information for "FAF1 cofactor enhances UFD1/NPL4-p97 unfolding efficiency across ubiquitin chain lengths independent of SUMO2"

**This PDF file includes:**

Figures S1 to S7
Table S1

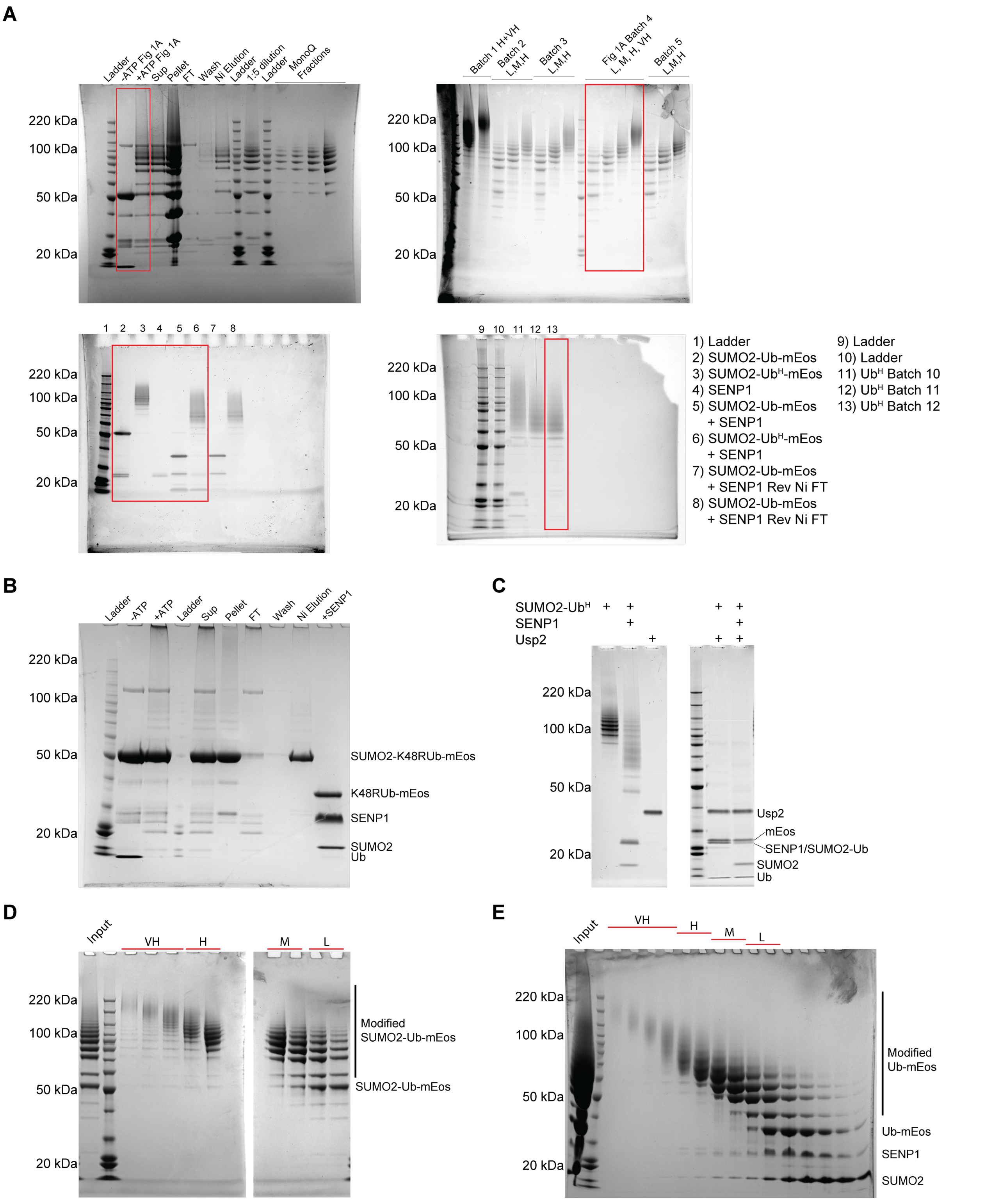


**Figure S1.** | **A)** Uncropped gels for Figures 1A and 2A. Red boxes indicate portion of gel utilized in main figure panels. **B)** *Substrates are modified with lysine-48 linked polyubiquitin chains.* A base substrate with K48R ubiquitin in the linear fusion was incubated with E1 *sp*Uba1, E2 *hs*UBE2K, and STUbL *sp*Slx8-Rfp2 and purified by nickel affinity (Ni-NTA) and treated with SENP1. Only minimal ubiquitination of the base substrate was observed and there is no ubiquitin chain on SUMO2. SYPRO-stained gel of reaction steps. **C)** Treatment of SUMO2-Ub$^{H}$ with Usp2 leads to collapse into SUMO2-Ub, mEos, and Ub bands. SYPRO-stained gel of reaction components. **D)** Coomassie-stained gel of fractions from size exclusion chromatography to purify SUMO2-Ub$^X$ substrates by size, pooled as indicated based on length of polyUb chain. **E)** Coomassie-stained gel of fractions from size exclusion chromatography to purify Ub$^X$ substrates by size and separate from SENP1 and SUMO2, pooled as indicated based on length of polyUb chain.

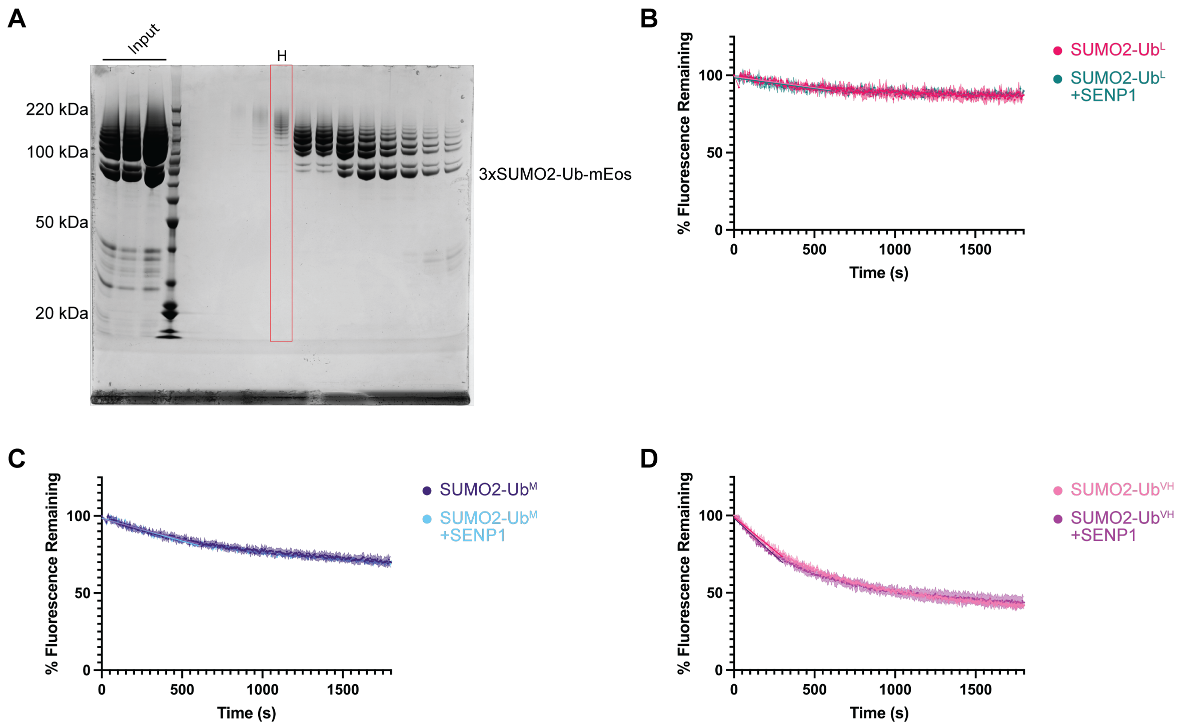


**Figure S2.** | **A)** Coomassie-stained gel of size exclusion chromatography fractions from 3xSUMO2-Ub$^{H}$-mEos substrate purification, red box indicates fraction used for assays. **B)** Unfolding SUMO2-Ub$^{L}$-mEos substrate +/- SENP1 by FUN-p97 with linear fits for first 600 seconds. **C)** Unfolding SUMO2-Ub$^{M}$-mEos substrate +/- SENP1 by FUN-p97 with linear fits for first 600 seconds. **D)** Unfolding SUMO2-Ub$^{VH}$-mEos substrate +/- SENP1 by FUN-p97 with linear fits for first 300 seconds

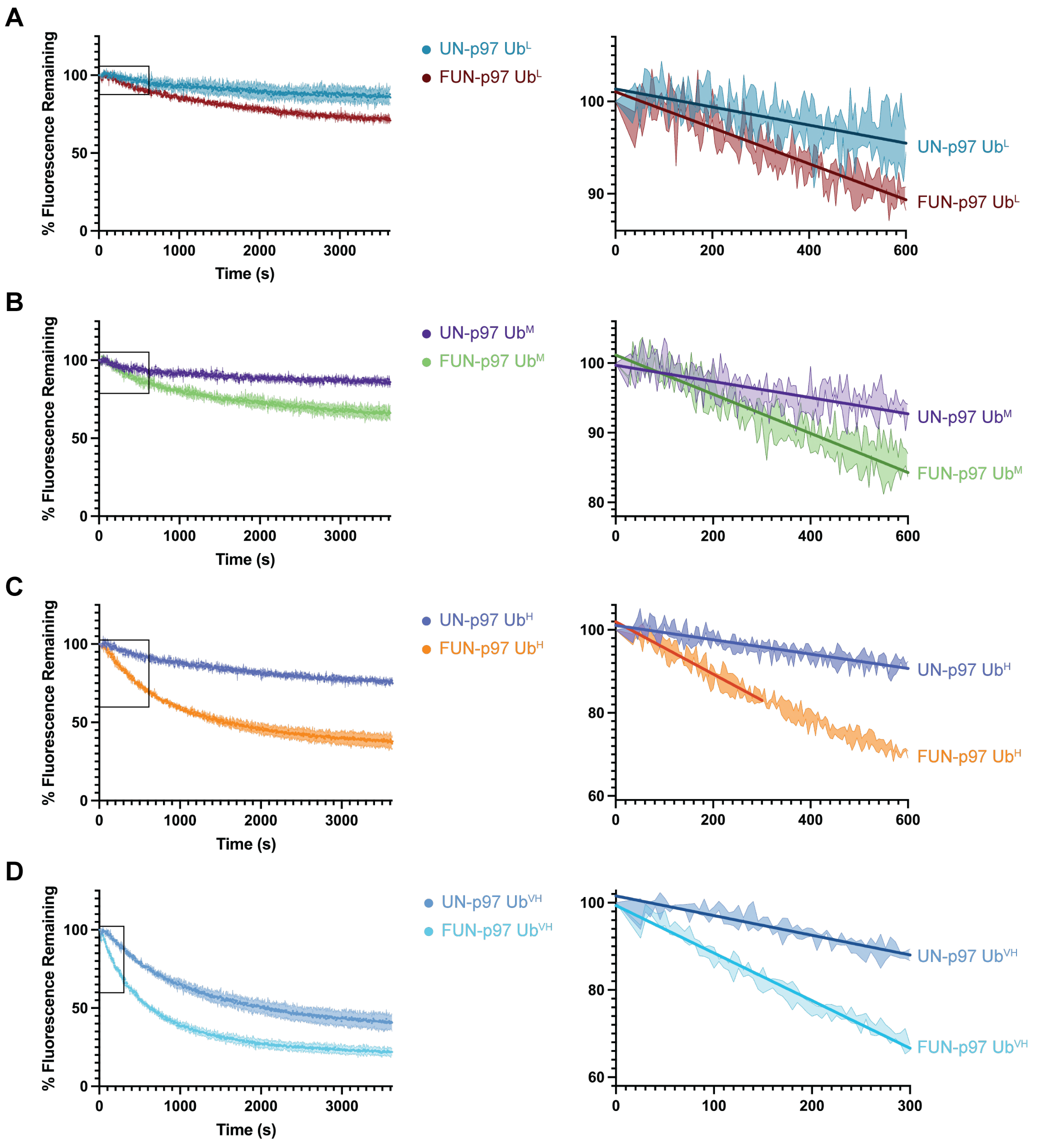


**Figure S3.** | **A)** Left: UN-p97 and FUN-p97 unfolding Ub$^{L}$-mEos. Right inset: first 600 seconds of unfolding as indicated by box on lefthand plot with linear fits. **B)** Left: UN-p97 and FUN-p97 unfolding Ub$^{M}$-mEos. Right inset: first 600 seconds of unfolding as indicated by box on lefthand plot with linear fits. **C)** Left: UN-p97 and FUN-p97 unfolding Ub$^{H}$-mEos. Right inset: first 600 seconds of unfolding as indicated by box on lefthand plot with linear fits. **D)** Left: UN-p97 and FUN-p97 unfolding Ub$^{VH}$-mEos. Right inset: first 300 seconds of unfolding as indicated by box on lefthand plot with linear fits.

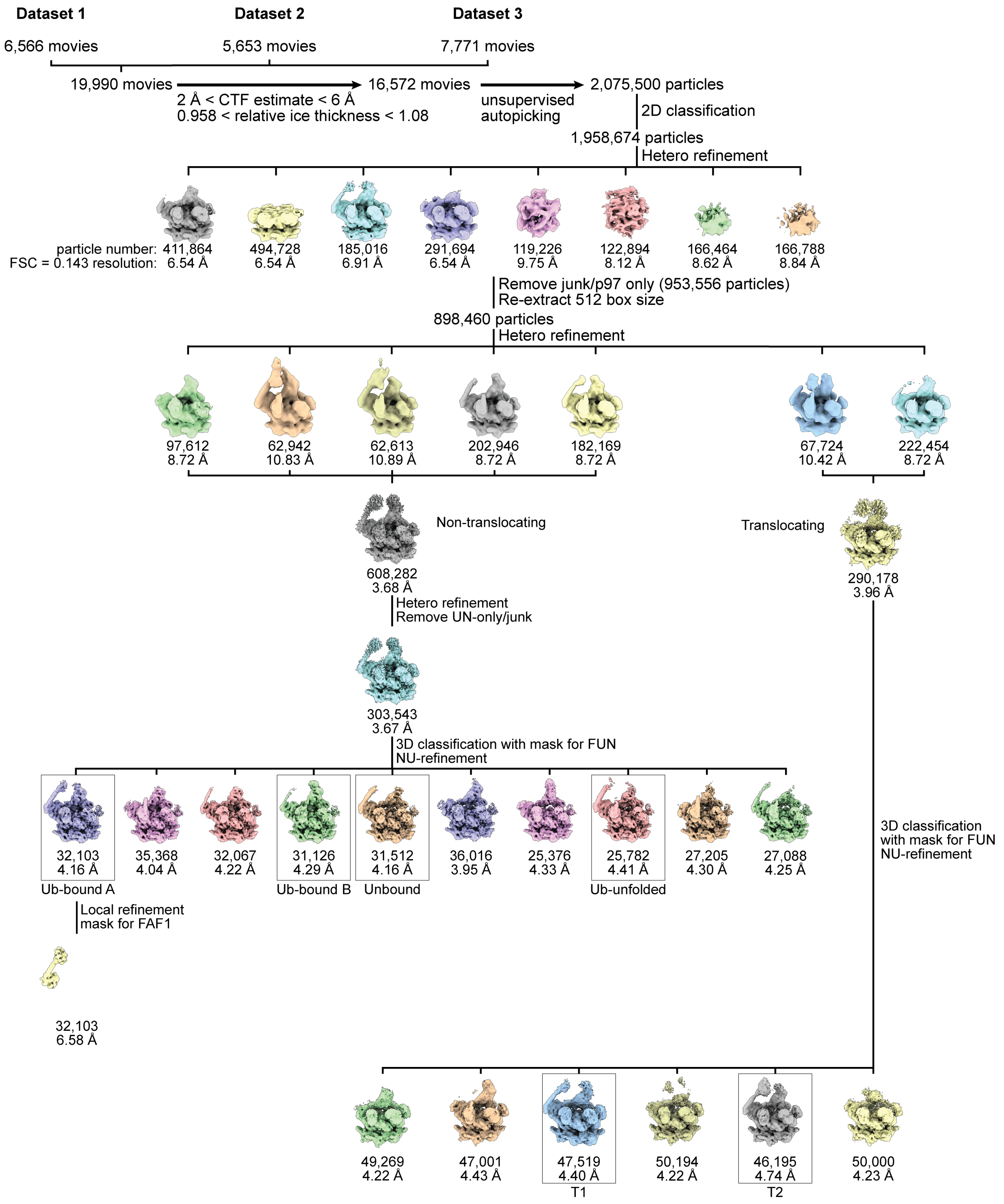


**Figure S4.** | **EM flowchart for data analysis**

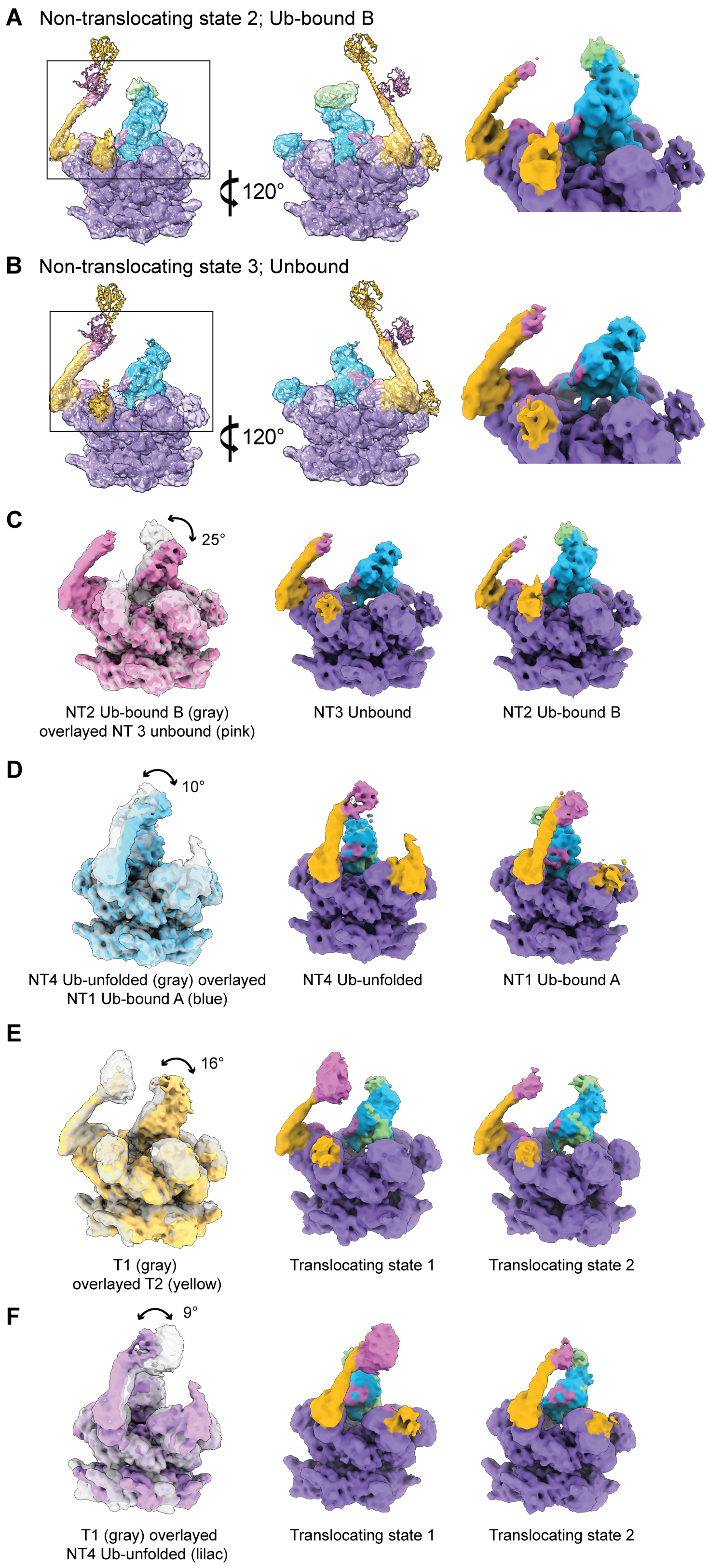
A Non-translocating state 2; Ub-bound B
120°
B Non-translocating state 3; Unbound
120°
C
25°
NT2 Ub-bound B (gray) overlayed NT 3 unbound (pink)
NT3 Unbound
NT2 Ub-bound B
D
10°
NT4 Ub-unfolded (gray) overlayed NT1 Ub-bound A (blue)
NT4 Ub-unfolded
NT1 Ub-bound A
E
16°
T1 (gray) overlayed T2 (yellow)
Translocating state 1
Translocating state 2
F
9°
T1 (gray) overlayed NT4 Ub-unfolded (lilac)
Translocating state 1
Translocating state 2

**Figure S5. |** Ribbon diagrams and density maps are colored as follow: FAF1 (yellow), UFD1 (pink), NPL4 (blue), p97 (purple), ubiquitin (green). **A)** *Non-translocating (NT) state 2 substrate-bound FUN-p97 low pass filtered to 6 Å; 'Ub-bound B'.* Left: AlphaFold model FAF1-UAS-helix-UBX, PDB 11TA (p97, UFD1, NPL4, FAF1 UBX, unfolded Ub), and PDB 1UBQ (folded Ub) rigid body fit into two views of electron density map. Right: Close-up view of interactions between NPL4, UFD1, FAF1, ubiquitin, and p97. **B)** *NT state 3 of FUN-p97 prior to substrate interaction low pass filtered to 6 Å; 'Unbound'.* Left: AlphaFold model FAF1-UAS-helix-UBX and PDB 11TA (p97, UFD1, NPL4, FAF1 UBX) rigid body fit into electron density map. Right: Close-up view of interactions between NPL4, UFD1, FAF1, and p97. **C)** Left: overlay of NT Ub-bound B (gray) and NT Unbound (pink) showing changing position of NPL4 prior to unfolding initiation. Angle of relative rotation marked, calculated using planes with a fulcrum at the center of the D1 pore. Middle: view of NT Unbound state. Right: view of NT Ub-bound B state. **D)** Left: overlay of NT Ub-unfolded (gray) and NT Ub-bound A (blue) showing different positioning of FAF1 during unfolding initiation. Angle of relative rotation marked, calculated using planes with a fulcrum at the FAF1 UBX domain. Middle: side view of NT Ub-unfolded state. Right: side view of NT Ub-bound A state. **E)** Left: overlay of translocating classes T1 (gray) and T2 (yellow) showing changing position of NPL4 during unfolding. Angle of relative rotation marked, calculated using planes with a fulcrum at the center of the D1 pore. Middle: view of T1 state. Right: view of T2 state. **F)** Left: overlay of T1 state (gray) and NT Ub-unfolded (lilac) showing different positioning of FAF1 during unfolding process. Angle of relative rotation marked, calculated using planes with a fulcrum at the FAF1 UBX domain. Middle: side view of T1 state. Right: side view of T2 state.

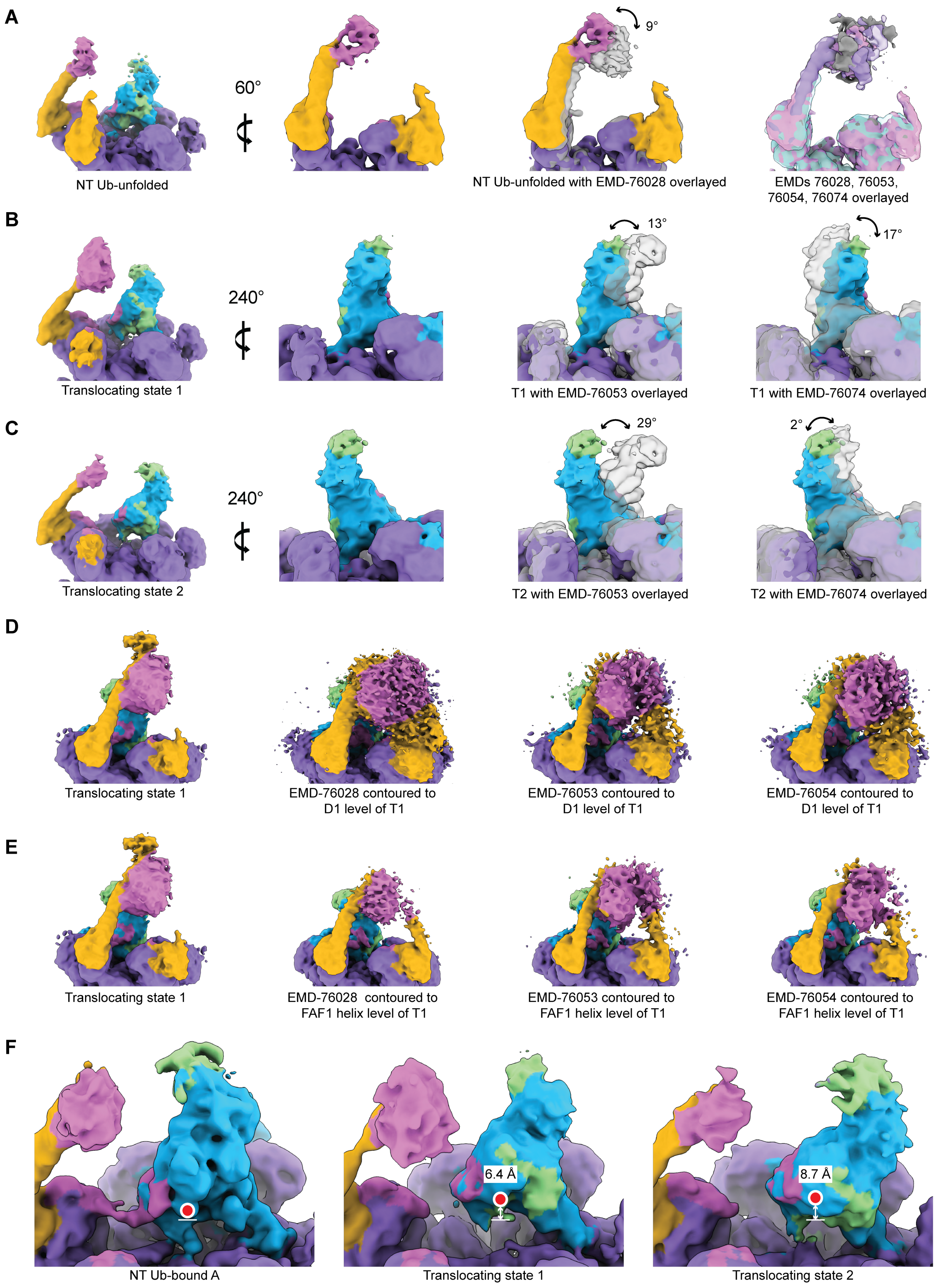
A
60°
9°
NT Ub-unfolded
NT Ub-unfolded with EMD-76028 overlayed
EMDs 76028, 76053, 76054, 76074 overlayed
B
240°
13°
17°
Translocating state 1
T1 with EMD-76053 overlayed
T1 with EMD-76074 overlayed
C
240°
29°
2°
Translocating state 2
T2 with EMD-76053 overlayed
T2 with EMD-76074 overlayed
D
Translocating state 1
EMD-76028 contoured to D1 level of T1
EMD-76053 contoured to D1 level of T1
EMD-76054 contoured to D1 level of T1
E
Translocating state 1
EMD-76028 contoured to FAF1 helix level of T1
EMD-76053 contoured to FAF1 helix level of T1
EMD-76054 contoured to FAF1 helix level of T1
F
6.4 Å
8.7 Å
NT Ub-bound A
Translocating state 1
Translocating state 2

**Figure S6. |** Density maps are colored as follow: FAF1 (yellow), UFD1 (pink), NPL4 (blue), p97 (purple), ubiquitin (green). **A)** *FAF1 position in NT Ub-unfolded differs from other FUN-p97 reconstructions.* Left: NT Ub-unfolded reconstruction. Left inset: view of FAF1 in NT Ub-unfolded. Middle inset: NT Ub-unfolded overlaid with EMD-76028. Angle of relative rotation marked, calculated using planes with a fulcrum at the FAF1 UBX domain. Right inset: EMDs 76028, 76053, 76054, and 76074 overlaid to show same approximate FAF1 angle in each reconstruction. **B)** *NPL4 position in Translocating state 1 differs from NPL4 in other FUN-p97 reconstructions.* Left: T1 reconstruction. Left inset: view of NPL4 in T1 reconstruction. Middle inset: T1 reconstruction overlaid with EMD-76053. Right inset: T1 reconstruction overlaid with EMD-76074. Angle of relative rotations marked, calculated using planes with a fulcrum at the center of the D1 pore. **C)** *NPL4 position in Translocating state 2 differs from NPL4 in other FUN-p97 reconstructions.* Left: T2 reconstruction. Left inset: view of NPL4 in T2 reconstruction. Middle inset: T2 reconstruction overlaid with EMD-76053. Right inset: T2 reconstruction overlaid with EMD-76074. Angle of relative rotations marked, calculated using planes with a fulcrum at the center of the D1 pore. **D)** *Translocating state 1 has additional density which may represent FAF1 UAS domain.* Left: reconstruction of T1 state. Second from left: EMD-76028 contoured to comparable level of D1 of T1 state. Second from right: EMD-76053 contoured to comparable level of D1 of T1 state. Right: EMD-76054 contoured to comparable level of D1 of T1 state. **E)** *Translocating state 1 has additional density which may represent FAF1 UAS domain.* Left: reconstruction of T1 state. Second from left: EMD-76028 contoured to comparable level of FAF1 helix of T1 state. Second from right: EMD-76053 contoured to comparable level of FAF1 helix of T1 state. Right: EMD-76054 contoured to comparable level of FAF1 helix of T1 state. **F)** *NPL4 moves away from p97 during translocation.* Left: NT Ub-bound A reconstruction. Middle: T2 reconstruction. Right: T1 reconstruction. Upward motion of NPL4 indicated for T1 and T2. Displacement calculated using position of NPL4 Gln 390 in rigid body fit model. Approximate location of Gln 390 in each state is marked with a red dot and position of Ub-bound A Gln 390 is marked with a white line with displacements indicated by white arrows.

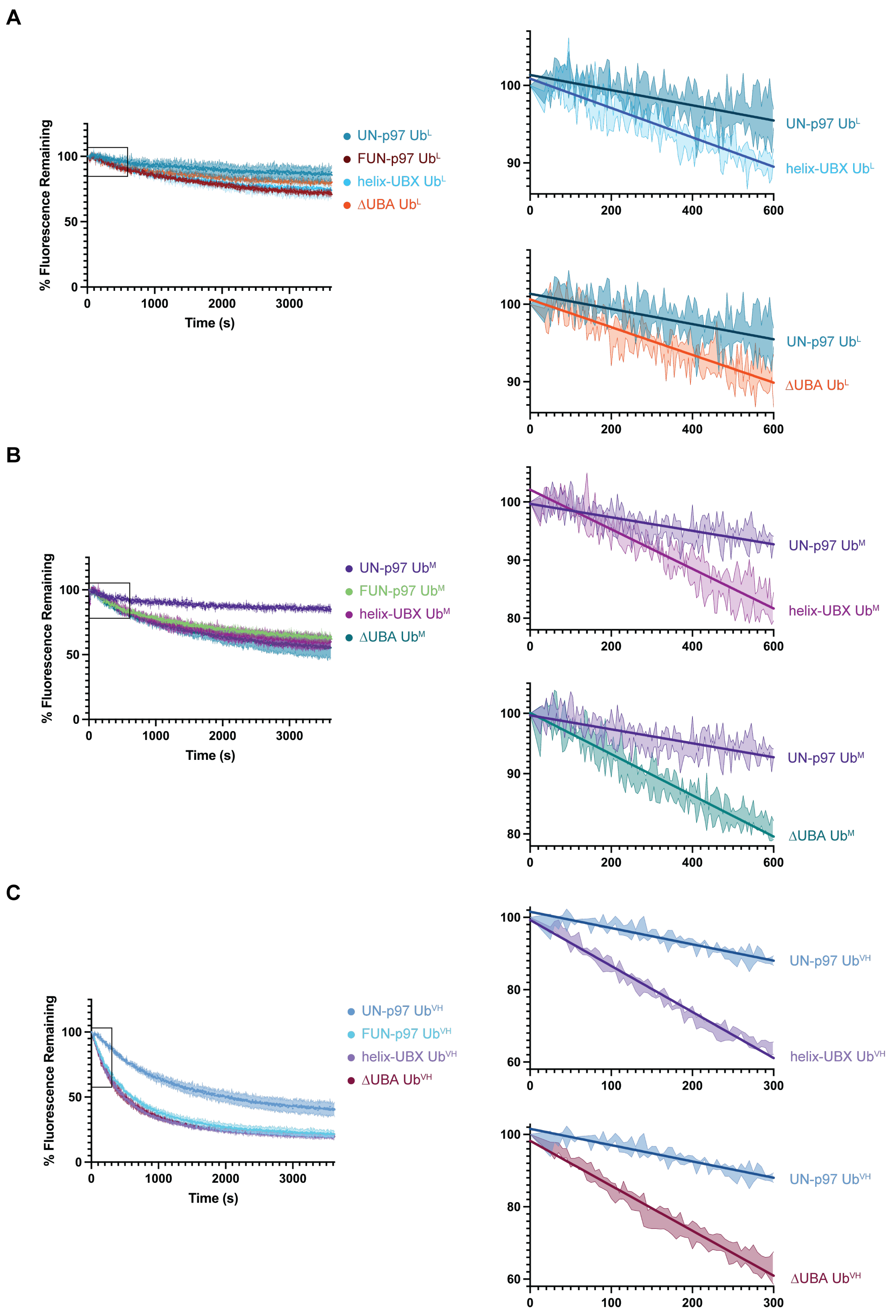


**Figure S7.** | **A)** Left: Unfolding $Ub^L$-mEos substrate by UN-p97, FL FUN-p97, or mutant FUN-p97 with helix-UBX or ΔUBA FAF1 constructs. Top right: first 600 seconds of UN-p97 and helix-UBX unfolding as indicated by box on lefthand plot with linear fits. Bottom right: first 600 seconds of UN-p97 and ΔUBA unfolding as indicated by box on lefthand plot with linear fits. **B)** Left: Unfolding $Ub^{M}$-mEos substrate by UN-p97, FL FUN-p97, or mutant FUN-p97 with helix-UBX or ΔUBA FAF1 constructs. Top right: first 600 seconds of UN-p97 and helix-UBX unfolding as indicated by box on lefthand plot with linear fits. Bottom right: first 600 seconds of UN-p97 and ΔUBA unfolding as indicated by box on lefthand plot with linear fits. **C)** Left: Unfolding $Ub^{VH}$-mEos substrate by UN-p97, FL FUN-p97, or mutant FUN-p97 with helix-UBX or ΔUBA FAF1 constructs. Top right: first 300 seconds of UN-p97 and helix-UBX unfolding as indicated by box on lefthand plot with linear fits. Bottom right: first 300 seconds of UN-p97 and ΔUBA unfolding as indicated by box on lefthand plot with linear fits. Data from Fig. S3 included for comparison between unfolding by UN-97, FUN-p97 with full-length FAF1, and FUN-p97 with truncated FAF1 constructs.

### Table S1. |. EM data and refinement

| | FUN-p97 bound to ubiquitinated substrate | | | | | |
|---|---|---|---|---|---|---|
| **Data collection** | Dataset #1 | | Dataset #2 | | Dataset #3 | |
| Microscope | Titan Krios G2 | | Titan Krios G2 | | Titan Krios G2 | |
| Detector/Mode | Gatan K3 Summit | | Gatan K3 Summit | | Gatan K3 Summit | |
| Mode | Counting - Super Resolution | | Counting - Super Resolution | | Counting - Super Resolution | |
| Data collection software | Serial EM | | Serial EM | | Serial EM | |
| Energy Filter | n/a | | n/a | | n/a | |
| Magnification | 22,500x | | 22,500x | | 22,500x | |
| Voltage (kV) | 300 | | 300 | | 300 | |
| Electron exposure ($e^-/Å^2$) | 70.7 | | 70.7 | | 70.7 | |
| Frames (collected/used) | 40/40 | | 40/40 | | 40/40 | |
| Exposure Time (s) | 3.995 | | 3.995 | | 3.995 | |
| Defocus range (μm) | -0.8 to -2.3 | | -0.8 to -2.3 | | -0.8 to -2.3 | |
| Super-resolution pixel size (Å) | 0.532 | | 0.532 | | 0.532 | |
| Fourier cropped pixel size (Å) | 1.064 | | 1.064 | | 1.064 | |
| Movies (collected/used) | 6566/4960 | | 5653/4434 | | 7771/7178 | |
| Initial particle projections (#) | 828,441 | | 826,664 | | 420,395 | |
| **Reconstructions** | Non-translocating state 1 Ub-bound A | Non-translocating state 2 Ub-bound B | Non-translocating state 3 Unbound | Non-translocating state 4 Ub-unfolded | Translocating state 1 | Translocating state 2 |
| Final particle projections (#) | 32,103 | 31,126 | 31,512 | 25,782 | 46,195 | 47,519 |
| Symmetry | C1 | C1 | C1 | C1 | C1 | C1 |
| Map resolution (Å) FSC threshold = 0.143 | 4.16 (6 low pass) | 4.29 (6 low pass) | 4.16 (6 low pass) | 4.41 (6 low pass) | 4.74 (6 low pass) | 4.40 (6 low pass) |
| Map resolution range (Å) Box (contoured) | 3.17-13.32 (3.7-11.44; 0.11 sig) | 3.20-14.37 (3.72-10.18; 0.105 sig) | 3.19-13.62 (3.76-10.01; 0.11 sig) | 3.42-14.16 (3.78-9.68; 0.1 sig) | 3.47-15.02 (3.59-12.38; 0.085 sig) | 3.41-14.28 (3.86-11.82; 0.085 sig) |
| Map sharpening B factor ($Å^2$) | -60.5 | -66.7 | -60.7 | -64.6 | -127.9 | -97.6 |
| Sphericity (FSC threshold = 0.5) | 0.835 | 0.881 | 0.792 | 0.924 | 0.909 | 0.871 |
| EMDB | 78389 | 78396 | 78395 | 78390 | 78397 | 78391 |
